# Multiplexed mapping of long-range projections with FluoCode, a combinatorial fluorescent barcoding system

**DOI:** 10.64898/2026.11.08.765127

**Authors:** Daichi Moriyasu, Akiya Watakabe, Itaru Imayoshi, Satoshi Fujimoto, Biswanath Saha, Yuki Ishida, Hikari Takeshima, Mizuki Sakamoto, Henrik Skibbe, Junichi Hata, Hinako Oshiro, Hideyuki Okano, Takeshi Imai

## Abstract

Comprehensive mapping of long-range projections remains a major challenge in neuroscience. Conventional virus tracing labels only one or a few pathways per animal, and inter-individual comparison requires registering whole-brain images to a standard atlas. However, inter-individual variability in morphology and wiring limits fine-scale mapping. Here we report “FluoCode”, a combinatorial fluorescent barcoding system for multiplexed projection mapping within single animals. We developed adeno-associated virus (AAV) vectors that express two of seven different fluorescent proteins, yielding 21 distinct color codes, and a “FluoCode Reader” that automatically classifies neurite signals by color combinations and morphology. Using FluoCode, we mapped projections from up to 21 cortical areas to subcortical regions in individual mice, revealing fine-scale topographic maps in small subcortical nuclei. In the marmoset, mapping from the primary visual cortex demonstrated topographic projections in higher visual areas and thalamic nuclei. FluoCode thus enables fine-scale, within-animal comparison of long-range projections in rodents and primates.

## INTRODUCTION

Neuronal wiring specificity defines many brain functions. Comprehensive mapping of neuronal connectivity across the entire nervous system is an ongoing challenge in neuroscience ^1,2^. Each neuron communicates with thousands of neurons through both local and long-range connections. Volume electron microscopy (vEM) is the gold standard in synaptic-resolution connectomics and can now reveal very detailed maps of local connectivity at the scale of cortical columns and areas ^3–5^. Comprehensive maps of long-range projections (projectome) have generally been acquired using light microscopy. Adeno-associated virus (AAV) vectors expressing fluorescent proteins (XFPs) are injected into one or a few brain regions, and whole-brain fluorescence images are acquired ^6^. The whole-brain images are then deformed and registered to a reference brain atlas based on cytoarchitectural landmarks ^7^. Based on the standardized platform of injection experiments, the mesoscale projectome data have been acquired for hundreds of brain regions in mice ^8^. This dataset now serves as an important resource for the neuroscience community ^8–10^. Single-neuron reconstruction, based on sparse single-color labeling, has revealed more detailed connectivity data for different cell types in both mice and primates ^11–16^.

While current approaches to describing the mesoscale projectome are important for establishing a reference connectivity map of the wild-type brain, they have several limitations. First, although the cytoarchitecture of the brain is commonly used to define landmarks for aligning multiple brain samples, the lack of precise, circuit-level landmarks poses a challenge to this process ^17–22^. Unlike genomic sequences, which can be aligned to a reference genome with single-nucleotide precision, brains lack intrinsic coordinates for such precise alignment. Furthermore, although overall connectivity is conserved among individuals, considerable morphological and wiring variability exists among them. Therefore, it is difficult to study fine-scale organization, such as topographic projections within small brain nuclei, wiring to regions with poorly defined cytoarchitecture, and connections with peripheral ganglia and organs, using approaches that compare multiple brain samples.

Second, neuronal wiring is not static but dynamic during development and under disease or injury conditions ^23,24^. Because every brain’s wiring is unique and evolves over the course of a lifetime, average maps cannot capture variability among individuals or changes across life stages. It is thus more informative to create wiring diagrams for multiple brain regions in individual brains than to merge multiple brain data into a common atlas. Third, the current approach, which uses many animals, cannot easily be applied to non-traditional model organisms, such as primates, due to their limited availability and ethical issues ^14,16,25–27^. These limitations could be overcome if many projection pathways were labeled and distinguished within a single brain.

Several approaches have been developed for multiplexed neuronal labeling. Multicolor labeling is straightforward, but the number of discriminable fluorescent proteins is limited. Stochastic and combinatorial multicolor labeling methods, such as Brainbow and Tetbow, have been used to map small-scale circuits ^28–32^. However, as the number of labeled neurons increases, the same or similar color combinations appear in multiple neurons, a phenomenon known as the birthday problem, making accurate readout difficult ^31–33^. RNA barcode-based strategies such as MAPseq and BARseq have achieved analyses of projections from thousands of neurons using bulk and *in situ* sequencing, respectively ^34,35^. However, the spatial resolution and the sample volume are each limited. Currently, none of these methods can easily be extended to projectome mapping at the whole-brain scale.

Here, we report FluoCode, a combinatorial fluorescent barcoding system for multiplexed mapping of long-range projections within individual animals. FluoCode AAV vectors express two of seven spectrally distinct fluorescent proteins, generating 21 discrete color codes. Because each FluoCode labels neurons in distinct brain regions, we can avoid the birthday problem. Unlike nucleotide-based barcoding strategies, the FluoCode can be directly decoded from fluorescence images. By injecting multiple FluoCode vectors into different brain regions of a single animal, projections can be traced and compared within the same brain. We also developed the “FluoCode Reader,” a program that identifies neurites labeled with specific FluoCodes based on color and morphology, enabling automated analysis of large image datasets. Using FluoCode, we mapped the projections from up to 21 cortical areas to subcortical regions in individual mice, revealing topographic maps in small thalamic nuclei and the brainstem. We also revealed the fine-scale topographic maps of projections from the primary visual cortex to the higher visual areas and thalamic nuclei in the marmoset.

## RESULTS

### FluoCode vectors for 21-plexed neuronal labeling

Multicolor labeling is a straightforward way to compare projections from different brain regions in a single animal. We have previously demonstrated that up to seven XFPs can be used and distinguished for neurite tracing. All of these XFPs are bright and stable enough, even at thin axon terminals. In this study, we used the following XFPs: mTagBFP2 (B), mTurquoise2 (T), and mAmetrine1.2 (A), excited with a 405 nm laser; mNeonGreen (N) and YPet (Y), excited with a 488 nm laser; mRuby3 (R) and Crimson (C) (or its equivalent, tdKatushka2), excited with a 552 nm laser ^31^. For *in vivo* experiments, we used Crimson instead of tdKatushka2 because Crimson is more resistant to aggregation when used as a fusion protein ^36^. Although fluorescent proteins excited by the same laser have overlapping fluorescence spectra, the fluorescence signals from each of the fluorescent proteins can be easily separated using linear unmixing (Figures S1A-C) ^31^.

To increase the number of discriminable labels further, we employed a combinatorial labeling strategy. We constructed vectors expressing two of the seven different XFPs, generating 21 (= _7_C_2_) color combinations, which we named FluoCodes (Figures 1A, B). We designated these FluoCodes with the two-letter abbreviations of the expressed fluorescent proteins (e.g., NR for mNeonGreen and mRuby3). FluoCodes are a type of protein-based barcoding technique that can be easily and accurately identified using fluorescence microscopy ^32,33,37^.

**Figure 1.**
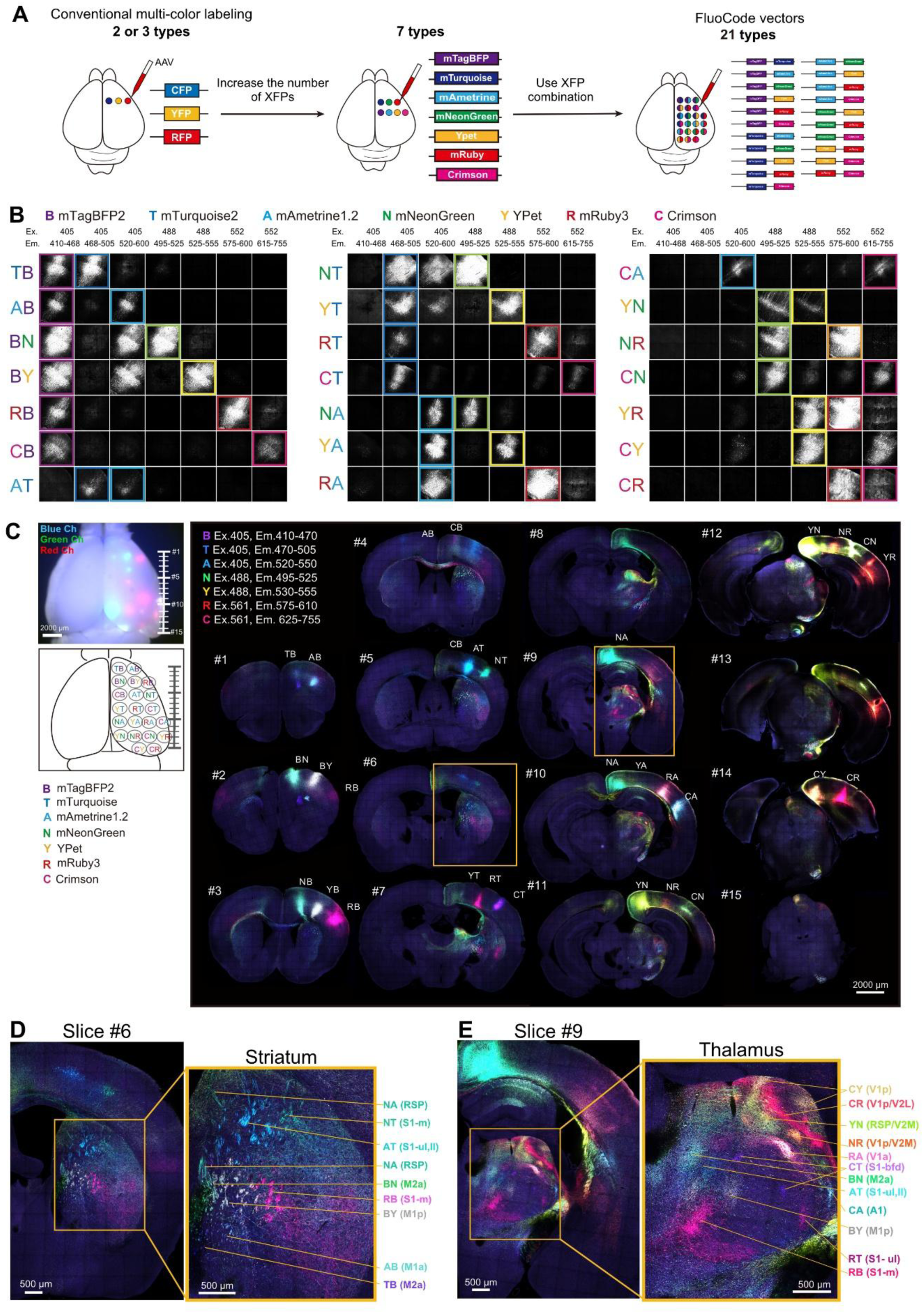
FluoCode: a combinatorial fluorescence barcoding strategy for projectome mapping. (A) Conceptual overview of the FluoCode system. Conventional multi-color AAV labeling uses two or three spectrally distinct fluorescent proteins (XFPs). Increasing the number of labels to seven increases the number of single-XFP labels to seven (middle), and expressing two of the seven XFPs within single cells yields 21 distinguishable FluoCodes (₇C₂ = 21; right). Each FluoCode is encoded by an AAV vector carrying two XFPs. The two XFPs are separated by an EV linker or a P2A. Abbreviations for XFPs are as follows: B, mTagBFP2; T, mTurquoise2; A, mAmetrine1.2; N, mNeonGreen; Y, YPet; R, mRuby3; and C, Crimson (or K, tdKatushka2). (B) Evaluation of the 21 FluoCode vectors expressed *in vivo*. One type of FluoCode AAV (TRE-XFP1-EVlinker-XFP2-WPRE) was co-injected with syn-tTA2 AAV into the cortex in each mouse (one FluoCode per mouse). Images of the injection site were obtained by confocal microscopy in seven channels and separated by linear unmixing. Each row shows one FluoCode, represented by the two-letter abbreviation of an XFP pair (e.g., TB for mTurquoise2 and mTagBFP2). The excitation (Ex) and emission (Em) wavelengths (in nm) of each channel are indicated at the top. Colored boxes highlight the two channels expected to contain the XFP signals. All 21 FluoCodes were detected in the expected channels in an all-or-none fashion, except for FRET-derived leakage into the mAmetrine1.2 channel in BN, BY, NT, and YT (See Figure S5). (C) A dorsal view of a whole mouse brain imaged in three spectral channels (blue, green, and red), showing the 21 injection sites arranged in a grid-like pattern in the right hemisphere (top). The anterior-posterior positions of the coronal sections shown in the right panel are indicated (#1–#15). The bottom left panel shows a schematic of the injection sites, with each site labeled by its two-letter abbreviation. Coronal sections (500 μm thick) spanning the entire brain from the anterior (#1) to the posterior (#15) regions, prepared using a vibratome and imaged with confocal microscopy (Leica TCS SP8) in seven channels (right). The excitation and emission wavelengths (Ex, Em, nm) used for each channel are indicated at the top left. The abbreviations above each section indicate the injection sites of the FluoCodes. Yellow boxes on sections #6 and #9 mark the regions shown at higher magnification in (D) and (E), respectively. The data presented here are from one animal out of three trials, with similar results. (D) Magnified images of the striatum in slice #6. Axonal projections from multiple cortical injection sites, distinguished by their FluoCodes after linear unmixing, form spatially segregated terminal fields within the striatum. Each labeled axon bundle is annotated with its FluoCode and the inferred cortical source area in parentheses (RSP, retrosplenial area; S1-m, primary somatosensory area, mouth; S1-ul, primary somatosensory area, upper limb; M2, secondary motor area; M1, primary motor area; S1-ll, primary somatosensory area, lower limb). A, anterior; P, posterior; M, medial; L, lateral. (E) Magnified images of the thalamus in slice #9, displayed as in (D). Corticothalamic axons from distinct cortical sites were largely non-overlapping within the thalamus. Annotations indicate the FluoCode and the inferred cortical source area (V1, primary visual area; S1-bfd, primary somatosensory area barrel field; V2M, secondary visual cortex, medial part; A1, primary auditory area).

When we expressed the 21 FluoCodes in HEK293T cells, many of them exhibited fluorescence in an all-or-none fashion following linear unmixing (Figure S1D). For *in vivo* applications, we expressed two XFPs as a fusion protein using a long, flexible linker (EV-linker) to ensure their uniform distribution within neurons ^38^. When each of the 21 FluoCode AAVs was injected into the mouse cortex (one FluoCode per mouse), fluorescence signals were detected in the expected two channels in an all-or-none fashion after linear unmixing (Figure 1B). Four FluoCodes (BN, BY, NT, and YT) showed leakage into the mAmetrine1.2 (A) channel due to FRET between the two XFPs. Nevertheless, we could easily identify which cells expressed which FluoCode (Figures S1E, F). Using a linear decoder based on the Euclidean distance across seven color dimensions, we accurately identified 99.7% of the cells expressing a FluoCode based on the fluorescence signals (Figure S1G).

### Visualizing projections from multiple cortical areas in a single mouse brain

We prepared adeno-associated virus (AAV) vectors that express each of the FluoCodes. To maximize the expression levels, we used a two-component system based on the Tet-Off system ^30,31^. Syn-tTA and TRE-FluoCode AAVs were co-infected. This system is also effective in minimizing trans-synaptic labeling. Because trans-synaptic infection is relatively inefficient, very few neurons express FluoCode due to the trans-synaptic spread of the two AAVs.

To unambiguously identify the FluoCode, it is important to ensure that each neuron expresses only one type of FluoCode. To avoid coinfection with multiple FluoCode AAVs, it is important to minimize overlap in the infection areas of different FluoCodes. We confined the expression of each FluoCode within a 1 mm diameter by optimizing the viral titer and the injection volume. We injected the FluoCode vectors into 21 cortical regions arranged in a grid pattern, with a minimum distance of 1.2 mm between regions. We then acquired 500-µm-thick brain slices spanning the entire brain and performed seven-channel confocal imaging (Figures 1C, S2A).

Our FluoCode vectors revealed the fine-scale topographic organization of projections from multiple cortical areas within a single brain. The axonal projections in the dorsal striatum (caudoputamen) maintain the spatial relationships of their cortical origins, confirming a preserved but graded topography (Figure 1D) ^10,39^. In the thalamus, projections from different cortical regions were discrete and segregated within each nucleus (Figure 1E), contrary to the graded map in the striatum. Together, these results demonstrate that our FluoCode vectors can resolve fine-scale projection patterns in multiple subcortical regions using just one mouse brain.

### Whole-brain imaging with block-face serial microscopy tomography (FASTd)

Next, we performed multiplexed mapping of corticofugal projections at the whole-brain scale in mice. The XFPs used in our FluoCodes can be excited by commonly used excitation lasers (405, 488, and 552 nm), which allows for versatile imaging applications. To visualize axonal projection profiles throughout the entire mouse brain, we used block-face serial microscopy tomography (FASTd), which is based on a spinning-disc confocal microscope (Dragonfly, Oxford Instruments) ^40,41^.

Due to the spectral constraints of the dichroic mirrors used in our Dragonfly system, we did not use FluoCodes with mTurquoise2 (T). We injected the remaining 15 FluoCode AAVs into dorsal cortical areas (Table S1) and performed whole-brain imaging using a 10× objective lens (NA 0.3) at 30 µm axial intervals (Figure 2A, Figure S2B, Video S1).

**Figure 2.**
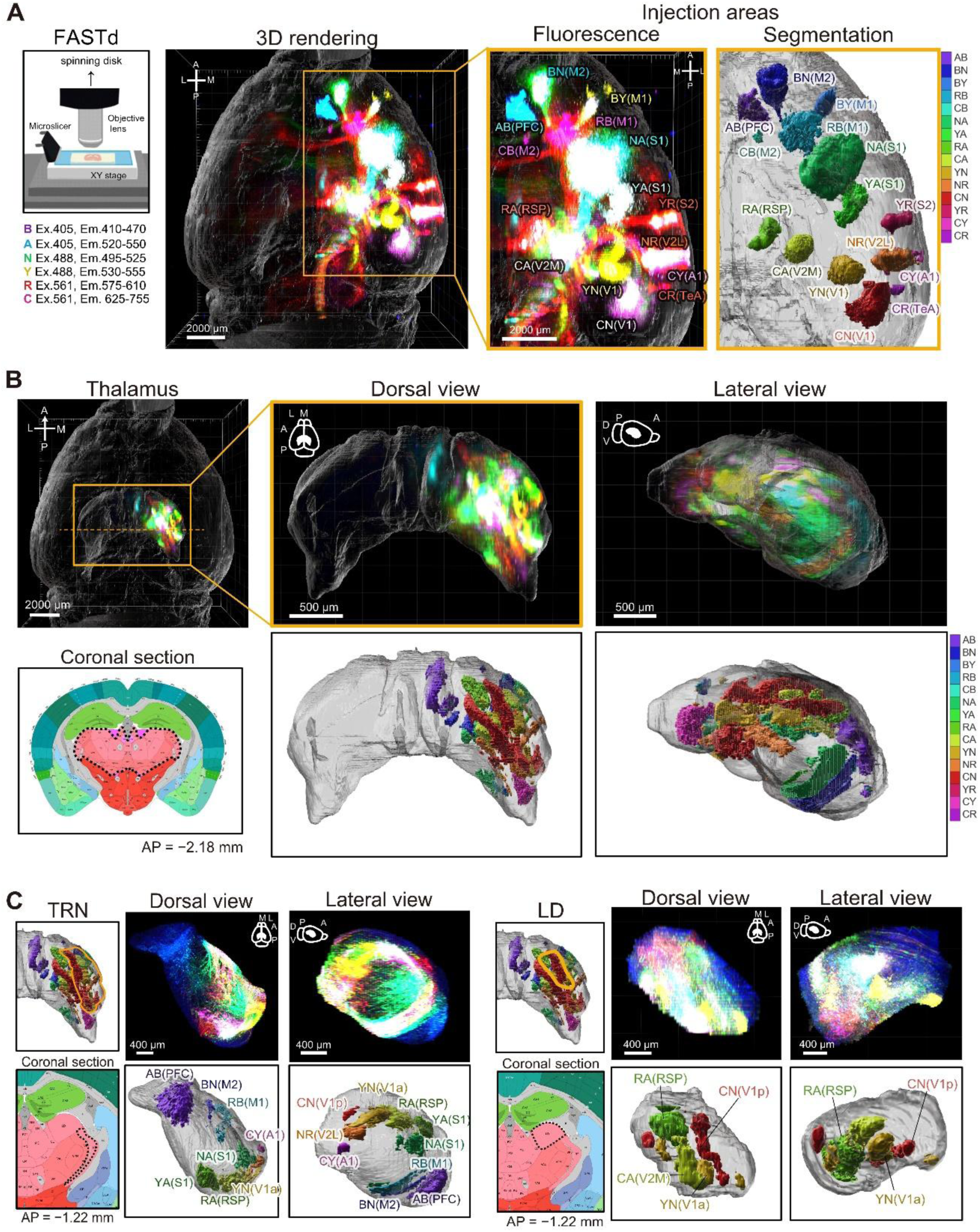
Whole-brain imaging of FluoCodes with FASTd in mice. (A) A schematic illustration of whole-brain imaging using FASTd (block-face serial microscopy tomography) with a 3D reconstruction of the mouse brain. The excitation and emission wavelengths (Ex, Em; nm) of the six channels are indicated below. FluoCode vectors were injected into 15 cortical areas in the right hemisphere. The right panel is a magnified view of the boxed region showing FluoCodes at the injection sites. The two-letter abbreviations of FluoCodes and injection areas are shown. The data presented here are from one animal out of two trials, with similar results. A, anterior; P, posterior; M, medial; L, lateral. Scale bars, 2,000 μm. See also Videos S1 and S2. (B) 3D rendering of corticothalamic projections. Top: Fluorescence signals of FluoCodes in the thalamus. Dorsal and lateral views are shown. Bottom: Segmentation of FluoCodes found in the corticothalamic axon terminals in the thalamus. (C) Topographic organization of corticothalamic projections in the thalamic reticular nucleus (TRN; left) and the laterodorsal nucleus (LD; right). For each nucleus, fluorescence (top: dorsal and lateral views) and segmentation (bottom: registered to the Allen CCFv3 Atlas) are shown. Multimodal inputs are arrayed in a topographic manner in these nuclei. Scale bars, 400 μm. See also Video S3.

Using the FASTd imaging system, we visualized segregated projections in multiple subcortical regions. The cortico-thalamic projection profiles from individual cortical areas were overall consistent with previous reports ^8^. The cortical topography was largely preserved in the thalamus. Frontal areas (medial prefrontal cortex [mPFC] and motor areas) tended to project to the anteromedial thalamus, while visual and auditory areas tended to project to the posterolateral regions (Figures 2B, S3A, Video S2). Furthermore, the cortical topography was also preserved in thalamic nuclei for specific modalities. For example, higher-order thalamic nuclei, such as the lateral posterior nucleus (LP; higher-order visual), exhibited topographic representation of cortical maps within specific sensory modalities (Figure S3B).

Some thalamic nuclei are known to receive multimodal inputs from multiple cortical areas. The thalamic reticular nucleus (TRN), a thin layer of GABAergic neurons surrounding the dorsal thalamus, receives collaterals from corticothalamic and thalamocortical axons. The TRN gates thalamocortical transmission by inhibiting thalamic relay neurons, suggesting a role for attention to a specific sensory modality. We found that the TRN receives projections from all cortical areas that we tested, including motor, sensory, and association areas. Inputs from different cortical areas were topographically arrayed, forming segregated domains (Figure 2C, Video S3). It is possible that the segregated map in the TRN is the anatomical basis for modality-specific attention ^42^. The laterodorsal (LD) nucleus is a higher-order thalamic nucleus, receiving multimodal inputs to mediate spatial navigation. We found that LD mostly receives cortical inputs from the retrosplenial cortex (RSP) and the visual areas (V1 and V2) and that these inputs are spatially segregated into the anterior and posterior domains, respectively (Figure 2C, Video S3). The feedback from RSP and V1/V2 may have distinct roles in spatial navigation ^43^. We also observed the topographic organization in the superior colliculus and callosal fibers (Figures S3C, D).

### Organization of descending cortical projections revealed by light-sheet microscopy

Light-sheet microscopy combined with tissue clearing is the most efficient strategy for large-scale volumetric imaging of mammalian tissues and organs. Although the resolution of the images is typically not as good as that of confocal imaging, this method can easily be applied to whole brains and even the entire body of mice ^44,45^.

To demonstrate the utility of our FluoCodes in light-sheet microscopy, we performed a multiplexed mapping of descending cortical projections. Excitation at 405 nm is not useful in light-sheet microscopy due to stronger light scattering of the cleared tissues and the higher absorbance of clearing agents. We, therefore, excluded FluoCodes with XFPs that are excited at 405 nm, such as those with mTagBFP and mAmetrine. mTurquoise2 was excited at 445 nm. We injected the remaining ten FluoCode AAVs into ten cortical areas spanning the motor and somatosensory areas (M1, M2, and S1). Then, we cleared the entire brain and spinal cord using CUBIC-L/R (Figures 3A, B) ^46^. We acquired the tiled images of the entire brain and the spinal cord with light-sheet microscopy using 1.1× (NA 0.1) or 4× (NA 0.35) objective lenses (Figures 3A, S4A, Video S4) ^47^.

**Figure 3.**
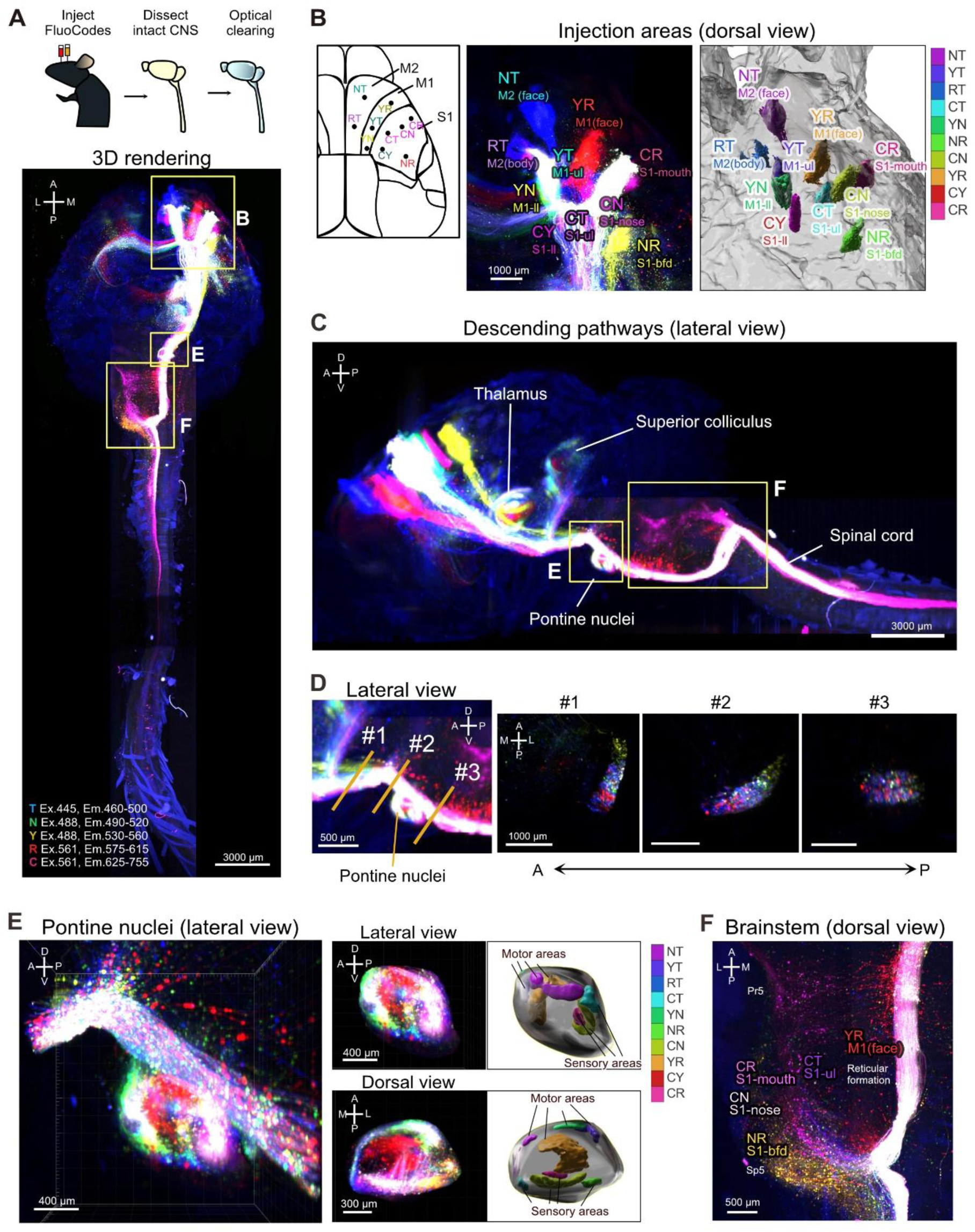
Multiplexed mapping of corticofugal projections in the mouse revealed with light-sheet microscopy. (A) Mapping of corticospinal projections using light-sheet microscopy with a 1.1× objective lens. FluoCode vectors were injected into the cortex. The brain and spinal cord were then dissected and cleared with CUBIC-L/R. The intact central nervous system was imaged by light-sheet microscopy (bottom, Sample #1). The yellow boxes indicate the regions shown in (B), (E), and (F). The data presented here are from one animal out of three trials, with similar results. Scale bar, 3,000 μm. See also Video S4. (B) Injection sites in the cortex. The schematic shows 10 injection sites distributed across motor and somatosensory areas. Raw fluorescence and segmentation of the injection sites are shown. The two-letter abbreviations of FluoCodes and injection areas are shown. ul, upper limb; ll, lower limb; bfd, barrel field. (C) A lateral view of the descending pathways. Axonal projections were found in the thalamus, superior colliculus, pontine nuclei, and the spinal cord. The yellow boxed region is shown in (E) and (F). D, dorsal; V, ventral; A, anterior; P, posterior. Scale bar, 3,000 μm. (D) Topography of corticopontine projections along the anteroposterior axis. Lateral view of the pontine nuclei. Cross-sections at the orange lines (#1–#3) are shown on the right. Scale bars, 1,000 μm. (E) Organization of corticopontine axon terminals. A lateral view of the pontine nuclei is shown on the left. Lateral (top) and dorsal (bottom) views of the terminal regions within the pontine nuclei are rendered on the right. Motor inputs terminate in the anterior and medial parts, while somatosensory inputs terminate in the central and posterior parts. Scale bars, 400 μm (lateral) and 300 μm (dorsal). (F) A dorsal view of the brainstem (boxed area in A). Axons from orofacial motor areas terminate in topographically distinct domains. Two-letter abbreviations of FluoCodes and injection areas are shown. Scale bar, 500 μm.

Multiple descending pathways from the labeled cortical areas were visualized in a single brain (Figure 3C). In the thalamus, cortical axons formed columnar terminal fields, which were similar to those observed with FASTd (Figure 3C). Axons from motor and somatosensory areas projected to the intermediate and deep layers of the superior colliculus (Figure 3C). In the cerebral peduncle, axon bundles from different cortical areas were topographically arranged down to the pontine nuclei, suggesting an axon sorting mechanism. Beyond that level, the axons with different FluoCodes became substantially intermingled (Figure 3D).

In the pontine nuclei, a relay station for the cortico-cerebellar network, both motor and somatosensory inputs were topographically segregated (Figure 3E). Motor areas projected to the anterior and medial regions, while somatosensory areas projected to the central and posterior regions, consistent with previous studies ^48–50^. The segregated map in the pontine nuclei may serve as the basis for divergent and convergent processing of copy signals from the cerebral cortex in the cerebellum.

Axons from the different orofacial areas terminated in a topographic manner in the brainstem, including the spinal trigeminal nucleus (Sp5), the principal sensory trigeminal nucleus (Pr5), and the reticular formation (Figure 3F) ^51^. Sp5 mainly relays the nociceptive inputs detected by trigeminal ganglion neurons. The topographic projection from S1 to these areas suggests precise feedback regulation at the early stage of somatosensory information processing. Axons from the forelimb and hindlimb areas of S1 and M1 descended through the pyramidal tract, crossing the midline at the pyramidal decussation (Figures 3F, S4A). These axons projected to the spinal cord and terminated in distinct domains of the contralateral spinal gray matter (Figure S4B). S1 axons were mostly found in the dorsal horn of the spinal cord, where somatosensory signals are processed and relayed ^52^. This result again suggests the top-down regulation of somatosensory signals. M1 axons were distributed in the intermediate zone, the ventral horn, and the medial dorsal horn (Figure S4C) ^53^. Together, our FluoCodes revealed the precise topographic organization of the descending cortical projections, not only for motor control but also for top-down modulation of somatosensory perception.

### FluoCode Reader for the automated identification of FluoCode signals

Although FluoCodes can be identified visually based on the combinatorial expression of XFPs, manual annotation and quantitative analysis are labor-intensive and nearly impossible for large image datasets. We, therefore, developed the “FluoCode Reader” program for the automated identification of the FluoCodes. To identify the FluoCodes in neurites with complex morphological features, our FluoCode Reader considers both color and morphological information (See **Methods** for details).

In the first stage of our FluoCode Reader, we calculated the images for each of the 21 FluoCodes (Step 1 in Figure 4A). For a FluoCode consisting of XFP-a and XFP-b, we first calculated the minimum intensity of these two XFP signals. Then, we subtracted the maximum intensity of the remaining XFPs. This eliminates pixels labeled with three or more XFPs and reduces contamination from autofluorescence and noise signals. For XFP pairs with possible Förster resonance energy transfer (FRET), namely BN, BY, NT, and YT, we made additional corrections at this stage (Figure S5). This yielded processed images for all the 21 FluoCodes.

**Figure 4.**
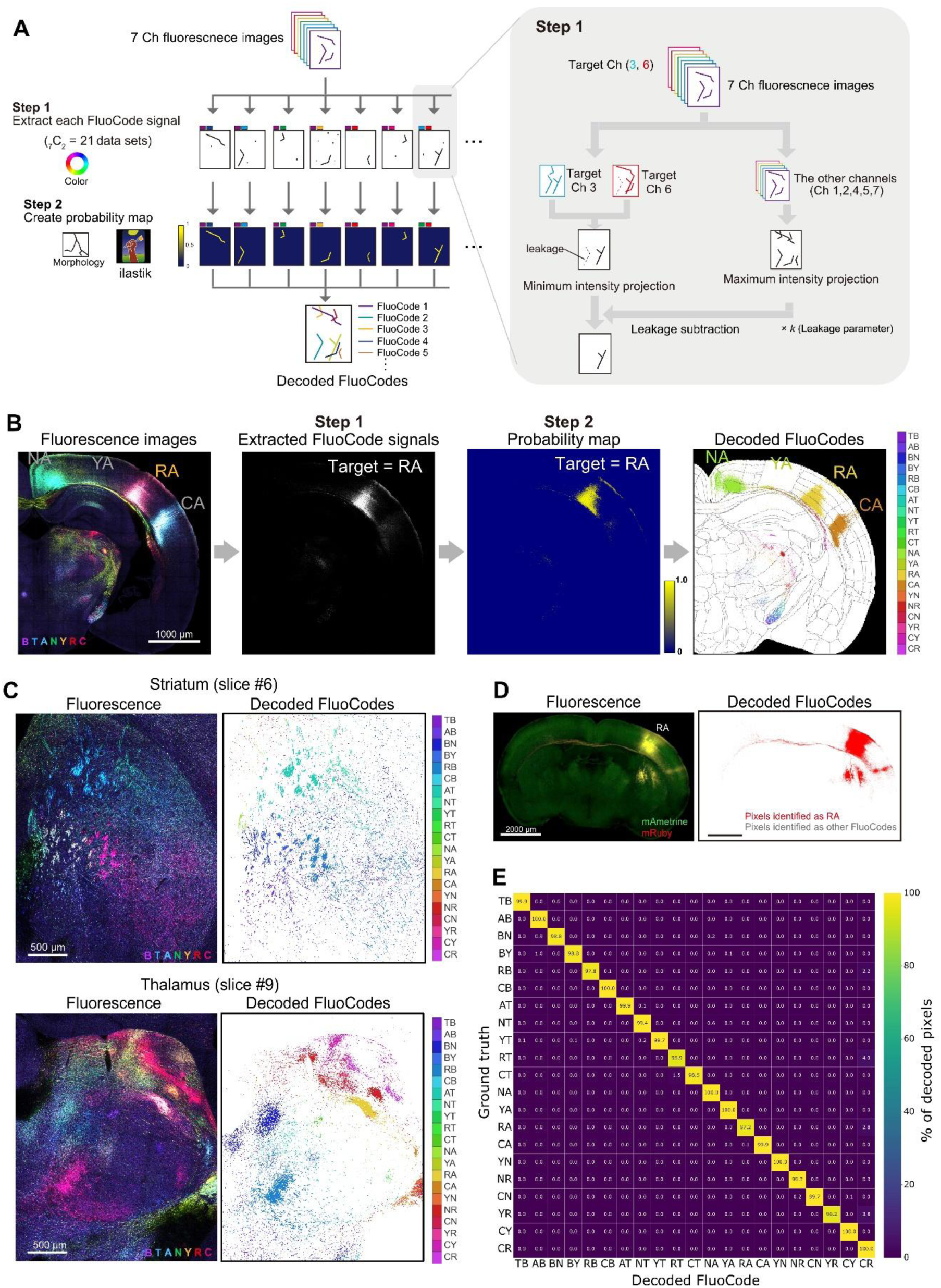
FluoCode Reader. (A) Overview of the FluoCode Reader pipeline. From the unmixed 7-channel image, the signal for each of the 21 FluoCodes is extracted (Step 1; see the inset on the right and Methods). Each FluoCode image was then converted into a probability map using ilastik (Step 2). Next, segmentation for FluoCodes was created, in which a single FluoCode with the highest and sufficient probability was chosen for each pixel. (B) An example of the image data processing for a mouse brain section labeled with FluoCodes (NA, YA, RA, and CA). From the unmixed 7-channel images, we generated extracted FluoCode signals for each target (Step 1). Ilastik probability maps for FluoCodes were then generated (Step 2). Decoded FluoCodes were overlaid on the Allen CCFv3 Atlas, where FluoCodes are color-coded. (C) Segmentation by the FluoCode Reader on a brain sample with 21 FluoCode injections into the cortex shown in Figure 1B. The FluoCode Reader was applied to the seven-channel confocal images of the mouse brain shown in Figure 1C-E. The raw fluorescence (left) and decoded FluoCode (right) images are shown for the striatum (slice #; top) and the thalamus (slice #9; bottom). (D) Neurite segmentation with FluoCode Reader. Raw fluorescence images and the segmentation are shown. Magnified views of the boxed areas are shown on the right. Only one type of FluoCode was expressed in each animal. The samples are the same as in Figure 1B. (E) Decoding performance of the FluoCode Reader. The rows indicate the injected (ground truth) FluoCode vectors, and columns indicate the FluoCodes decoded from the images using the FluoCode Reader. Values indicate the fraction of labeled pixels assigned to each FluoCode and were quantified from coronal sections expressing a single type of FluoCode. n = 1 mouse per FluoCode (21 mice in total). The data are from the images shown in (C) and Figure 1B.

Although the above process effectively extracts the signals for each FluoCode, the processed images still contain a lot of noise. In the second step, we used ilastik, an open source, machine learning-based segmentation tool, to extract specific morphological features, namely neurites and somata (Step 2 in Figure 4A) ^54^. Ilastik is based on a random forest algorithm and can easily be trained to detect specific morphological features using a limited training dataset. We trained the model to identify the neurite and soma signals for all the seven XFP channels (1-2 manually annotated images per channel). It only took one hour to prepare the training dataset, which can then be used to identify neurites in FluoCode images. For each of the 21 FluoCodes, ilastik calculates the probability maps for the neurite and soma signals. We assigned the FluoCode with the highest and sufficient probability to each pixel in the images based on the probability maps for all 21 FluoCodes. The final output is the decoded FluoCode images, in which the FluoCode ID (#1-21) is represented by a number in each pixel (Step 3 in Figure 4A).

In our proof-of-concept experiment, we applied the FluoCode Reader to unmixed seven-channel images of mouse brain samples, each of which expressed only one of 21 types of FluoCodes (Figure 4B, C). The FluoCode Reader successfully identified fine axonal fibers labeled with a specific FluoCode (Figure 4D). Importantly, there was minimal false-positive detection. On average, across the 21 FluoCodes, the FluoCode Reader assigned the correct (injected) FluoCode in 99.1% of labeled pixels (range, 95.9–100%). Fourteen of the 21 FluoCodes were identified with ≥99% accuracy (Figures 4D, E). Thus, the FluoCode Reader can accurately identify the neurites with specific FluoCodes based on seven-channel images (Figures 4B, C).

It should be noted that identifying the FluoCode is difficult when axons with different FluoCodes overlap in images. In brain samples with multiple FluoCode injections, we observed some false positives when axons with different FluoCodes were highly intermingled, especially in low-resolution images. The FluoCode Reader works best with high-resolution confocal images (≥10× objectives).

### The FluoCode Reader reveals a quantitative map of cortico-subcortical projections in a single mouse brain

We used the FluoCode Reader to describe the quantitative map of cortico-thalamic projections. We injected FluoCode AAVs into 15 cortical areas, obtaining high-resolution (20× objective) confocal images of consecutive brain slices spanning the entire thalamus (Figure 5A, Video S5). Using the FluoCode Reader, we automatically identified 14 FluoCodes in thalamic nuclei. As was seen in the FASTd data, axons from different cortical areas were segregated in most of the thalamic nuclei (Figures 5B, C, Video S5). This has been difficult to identify using conventional single-color labeling strategies ^9^. A quantitative cortico-subcortical projection map for the 14 cortical areas can be obtained from just one animal (Figures 5D, F).

**Figure 5.**
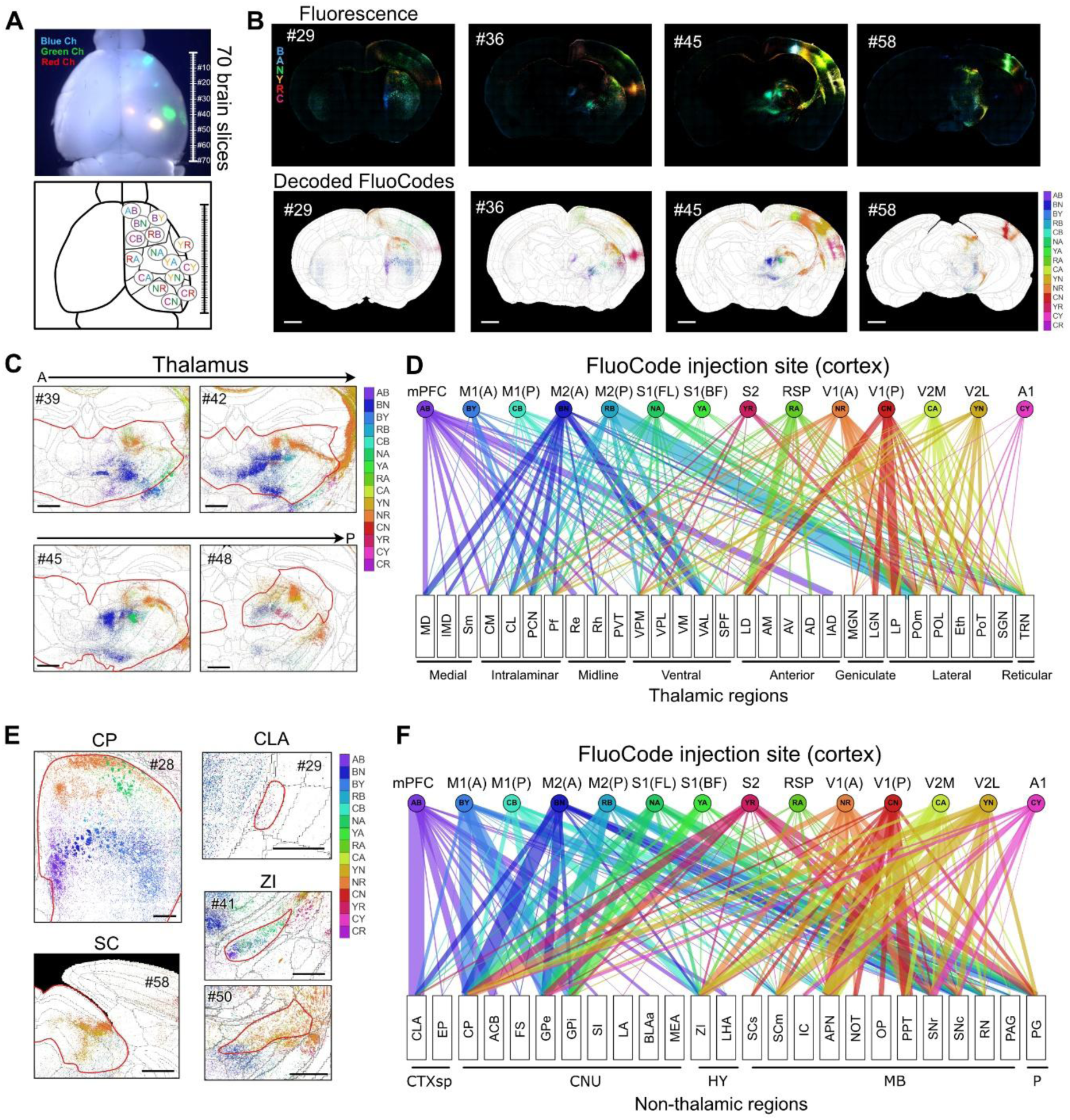
Quantitative map of cortico-thalamic projections in mice revealed with FluoCode Reader. (A) FluoCode vectors were injected into 15 cortical areas. This is a dorsal view of a whole mouse brain imaged with three spectral channels (blue, green, and red), showing the 15 injection sites in the right hemisphere. Serial coronal sections (70 sections, shown on the right) were analyzed with the FluoCode Reader. The data presented here are from one animal out of three trials, with similar results. See also Video S5. (B) Representative coronal sections (#29, #36, #45, and #58) before and after segmentation with FluoCode Reader. The segmentation images were overlaid onto the Allen CCFv3 Atlas using BigWarp. Scale bar, 1,000 μm. (C) Decoded FluoCodes in four representative sections spanning the thalamus (#39, #42, #45, #48; anterior-posterior). Red outlines delineate the thalamus. Axons from different cortical areas terminate in non-overlapping territories in most thalamic nuclei. Scale bar, 500 μm. (D) Quantitative map of corticothalamic projections obtained from a single animal. The width of the band represents the projection density. Thalamic nuclei are classified into eight groups: medial, intralaminar, midline, ventral, anterior, geniculate, lateral, and reticular. CR was excluded from the analysis because its expression level was low in this sample (see Methods). See Figure S6A for the quantitative data. (E) Decoded FluoCodes in representative non-thalamic regions: the caudoputamen (CP; section #28), claustrum (CLA; section #29), zona incerta (ZI; sections #41 and 50)), and superior colliculus (SC; section #58). The decoded FluoCode images were overlaid onto the Allen CCFv3 atlas. The red lines delineate each region. Scale bar, 500 μm. (F) Quantitative map of projections to non-thalamic subcortical regions. Target regions are classified into five groups: cortical subplate (CTXsp), cerebral nuclei (CNU), hypothalamus (HY), midbrain (MB), and pons (P). See Figure S6B for the quantitative data.

We observed both modality-specific and multimodal thalamic nuclei. The first-order sensory and motor nuclei received projections from a limited set of functionally related cortical areas. S1 projected to the ventral posteromedial (VPM) nucleus and the ventral posterolateral (VPL) nucleus. Visual and auditory areas projected to the dorsal lateral geniculate (LGd) nucleus and the medial geniculate nucleus (MGN), respectively. Primary and secondary motor areas (M1 and M2) projected to the ventral nuclei, including the ventral anterior-lateral complex (VAL) and the ventromedial nucleus (VM). In contrast, higher-order nuclei received multimodal inputs from multiple cortical areas. The mediodorsal nucleus (MD) and the intralaminar nuclei (the central lateral nucleus, CL, and the central medial nucleus, CM) received inputs predominantly from mPFC and M2. The posterior medial nucleus (POm) received inputs from both S1 and motor areas. Both the lateral posterior (LP) and laterodorsal (LD) nuclei received inputs from multiple visual areas, but the LD nucleus received denser projections from the retrosplenial cortex (RSP) (Figures 5D, S6A).

Similarly, we observed multimodal inputs in some non-thalamic subcortical regions (Figures 5E, F, S6B). For example, all of the injected cortical areas projected to the caudoputamen (CP) of the striatum, consistent with previous studies ^10^. The zona incerta (ZI) is also known to receive multimodal inputs to control behavioral and physiological state modulation ^55^. Indeed, we observed that the ZI receives projections from multiple cortical areas, but these inputs are segregated within the ZI (Figures 5E, F). The claustrum (CLA) is known to project to most cortical areas ^56^. However, the CLA received inputs predominantly from the medial prefrontal cortex (mPFC), consistent with previous reports.

To quantitatively compare our results with those of the Allen Mouse Brain Connectivity Atlas ^8^, we selected Allen experiments with the closest injection sites and calculated the projection density in thalamic regions (Figures S6A, B). There were high correlations between FluoCode and Allen data for most pairs (mean *r* = 0.774; all *P* < 0.01), suggesting similar subcortical projection patterns between the datasets (Figure S6C). Of course, the true strength of FluoCode lies in our ability to visualize spatial organization within small brain nuclei.

### Visualizing topographic projections in the marmoset brain

The organization of the primate brain differs considerably from that of rodents ^2,57^. Whole-brain mapping of projections has been performed in some areas of the primate brain, including the marmoset and the macaque ^14,16,25–27^. However, comprehensive mapping of the projectome in primates remains challenging due to their larger size and limited availability. Additionally, variability among individuals is a more problematic issue in primates than in inbred mouse strains.

To demonstrate the power of our approach in primates, we tested FluoCode vectors to characterize the topographic projections from the primary visual cortex (V1) in the marmoset. The topographic organization of the visual system is well-established in primates, making it ideal for proof-of-concept experiments. We used serial two-photon tomography (STPT) to image the entire marmoset brain. STPT is based on serial block-face imaging. Sections retrieved during STPT imaging were then imaged with spinning-disc confocal microscopy for in-depth analysis using the FluoCode Reader. In our pilot experiment, we found that mTagBFP was quenched after the prolonged sample preparation and processing time (∼1 month) required for STPT imaging (Sample #2, Figure S7F) ^6^. Additionally, since the primate brain exhibits higher autofluorescence, we acquired two additional fluorescence channels to subtract autofluorescence using unmixing (Figure S7A; see **Methods**).

We injected ten FluoCode AAVs into the exposed part of the right V1 and five FluoCode AAVs into the buried part of the left V1, ensuring that the injections were non-overlapping (Figures 6A, S7B, C). First, we performed STPT imaging of the whole marmoset brain at 100 µm z-intervals (Videos S6, S7). Although our STPT can only acquire three fluorescence channels (cyan, yellow, and red), it clearly visualized all of the FluoCode signals. Projections from V1 were found in several well-known targets, including the secondary visual area (V2), the middle temporal area (V5/MT), the superior colliculus (Figure S7E), the lateral geniculate nucleus (LGN), and the pulvinar nucleus. Brain sections retrieved during STPT imaging were also imaged with a high-resolution spinning-disc confocal microscope (6-color imaging with a 20× objective) to identify the FluoCodes.

**Figure 6.**
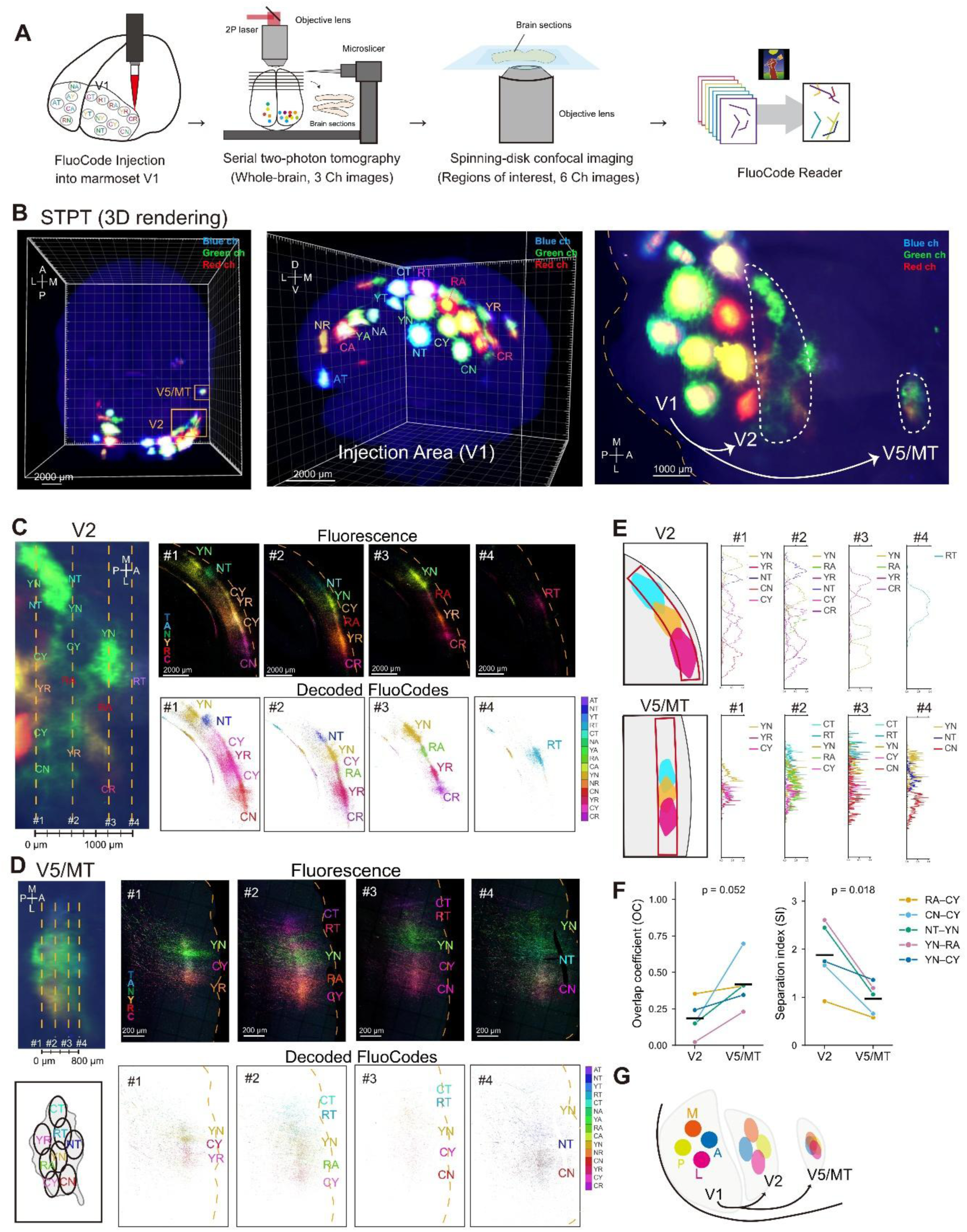
Topographic projections from V1 to higher visual areas in the marmoset. (A) Schematic illustration of the experimental pipeline. FluoCode vectors were injected into the primary visual cortex (V1) of the marmoset, followed by whole-brain, 3-channel imaging with serial two-photon tomography (STPT). Brain slices containing projection sites were subsequently imaged at a higher resolution with 6-channel spinning-disc confocal microscopy. FluoCodes were then identified with FluoCode Reader. (B) A 3D rendering of the entire marmoset brain imaged with STPT (Sample #1). FluoCode vectors were injected into five and ten sites in the left and right V1, respectively. Magnified views of the injection areas (caudal and lateral views) are also shown. Projections from V1 to V2 and V5/MT are visualized in the lateral view. Dotted lines indicate V2 and V5/MT (right). The two-letter abbreviations of the FluoCodes are shown. D, dorsal; V, ventral; M, medial; L, lateral; A, anterior; P, posterior. See also Videos S6, S7. (C) Spatial organization of the projections from V1 to V2. The positions of the sections (#1-4) are shown in the STPT images on the left. Fluorescence and segmentation images are shown for sections #1-4. (D) The spatial organization of the projections from V1 to V5/MT. (E) Quantitative maps of FluoCode signals in coronal sections spanning V2 (top) and V5/MT (bottom). (F) Quantification of axonal segregation in V2 versus V5/MT. The Overlap Coefficient (left) and Separation Index (right) calculated for five representative FluoCode pairs (RA-CY, CN-CY, NT-YN, YN-RA, and YN-CY). Lines indicate matched pairs between V2 and V5/MT. Each colored line connects the same FluoCode pair in the two areas. Black horizontal bars indicate means. (G) Summary of the topographic projections from V1 to higher visual areas. V5/MT shows higher overlap than V2.

First, we examined topographic projections from V1 to V2 and V5/MT areas (Figure 6B). Mirror-symmetric topographic projections were observed in both areas (Figures 6B-D), consistent with previous reports ^58^. Additionally, we found that the topography was more obscured in V5/MT than in V2 (Figures 6C-G). This result is consistent with established knowledge that the receptive field is broader in V5/MT neurons than in V2 neurons ^59^.

We also examined projections from V1 to the thalamus. We observed axonal projections in the LGN and the pulvinar nucleus, which are the first-and higher-order thalamic nuclei, respectively (Figures 7A, B). The pulvinar in primates is much more expanded and diverse than the LP in mice, suggesting that it plays a pivotal role in higher visual functions unique to primates ^60^. Unlike V2 and V5/MT, projections from each injection site were clearly segregated in the thalamus. In the LGN, each FluoCode exhibited a columnar structure that penetrated vertically through the layers of the LGN in mirror-symmetric positions (Figure 7C). Similar to the LGN, the signals in the pulvinar also exhibited columnar and topographic organization along the anterior-posterior axis (Figure 7D). Furthermore, we observed that the pulvinar has duplicate maps in mirror-symmetric positions: one in the dorsal region and the other in the ventral region. Projections from the posterior V1, which represents the central visual field, exhibited a clear spatial separation between the dorsal and ventral parts. In contrast, projections from the anterior V1, which represents the peripheral visual field, were found at the interface of the two maps (Figures 7E-G). The presence of duplicate pulvinar maps has also been suggested in macaque studies ^61,62^.

**Figure 7.**
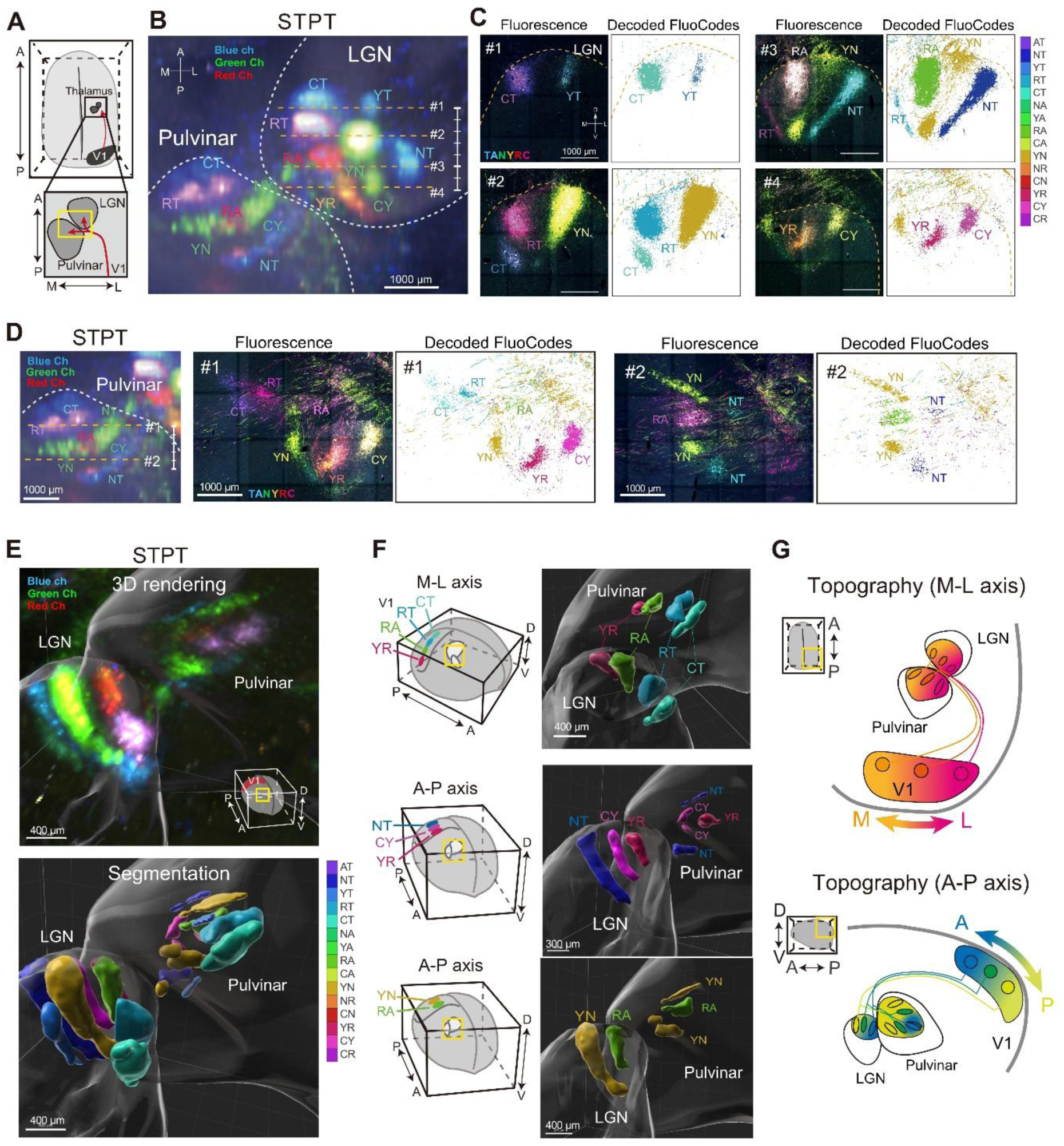
Organization of projections from V1 to thalamic nuclei in the marmoset. (A) Schematic of the thalamic projections from V1 (dorsal view). This is the same analysis shown in Figure 6. The enlarged area shows the lateral geniculate nucleus (LGN) and pulvinar. A, anterior; P, posterior; M, medial; L, lateral. (B) A representative STPT image showing fluorescence signals in the LGN and pulvinar (dorsal view). Yellow dotted lines indicate the locations of sections #1–4 in (C). White dotted lines delineate the LGN and pulvinar. (C) Topographic organization of V1 axons in the LGN. (D) Topographic organization of V1 axons in the pulvinar. The yellow dotted lines indicate the locations of sections #1, 2, which are shown in (E). (E) 3D rendering and segmentation of V1 axons in the thalamus. Segmentation was performed manually using Imaris based on the results of the FluoCode Reader. The LGN and pulvinar are rendered as semi-transparent surfaces. (F) The topographic organization of V1 projections to the thalamus. V1 axons are shown from an angle that best segregates different FluoCodes. (G) A summary of the topographic projections from V1 to the LGN and pulvinar. The ML and AP axes of the V1 are represented as distinct orthogonal axes in the thalamus. The pulvinar contains two mirror-symmetric maps.

Projections from each injection site were also topographically arranged in the superior colliculus (Figure S7E). Projections from the buried V1 (left hemisphere) were located in the more posterior superior colliculus and more anterior LGN compared to the projections from the exposed V1 (right hemisphere) (Figures S7D, E).

Together, these results demonstrate that our FluoCode vectors are powerful for investigating projections from multiple brain regions in primates.

## DISCUSSION

In this study, we developed FluoCode, a combinatorial fluorescent barcoding system that enables the mapping of up to 21 projection pathways within a single brain. Its coding capacity is much smaller than that of stochastic multicolor labeling, such as Brainbow/Tetbow, and nucleotide-based barcoding strategies, such as MAPseq and BARseq ^28–32,34,35^. However, because each FluoCode is assigned to the defined injection sites, FluoCode can avoid the birthday problem and enable highly accurate decoding based on fluorescence images (Figure 4). Therefore, FluoCode is complementary to existing approaches. Notably, our tool is compatible with various kinds of fluorescence microscopy systems, including confocal, spinning-disc confocal, and light-sheet microscopy, which use commonly available laser lines (405/445, 488, and 552 nm). The unmixing and FluoCode Reader programs are available as ImageJ plugins on GitHub. Thus, our new tool is accessible to many neuroscience laboratories using standard fluorescence microscopy systems.

Previous whole-brain imaging approaches required image deformation and registration to a standard atlas for multi-animal comparisons. However, every brain is different in terms of morphology and microscopic wiring diagrams. For example, neuronal locations and connectomes differ considerably among individuals of the same genetic background even in *C. elegans* ^24,63–66^. Furthermore, cytoarchitectural landmarks are insufficiently reliable to align brain circuits from different animals. Our multiplexed labeling approach is easy yet effective for studying fine-scale connectivity and/or variations among individuals. For example, our method may be useful for studying changes in the mesoscopic projectome in disease and injury models ^23^. In this study, we mapped neuronal populations using dense labeling. However, it should be possible to adjust the sparseness of the labeling in our two-component system to enable single-neuron tracing. Combining our method with cell type-specific mini-promoters in AAV may also be useful for cell type-specific mapping of the projectome ^67–69^.

FluoCode is particularly useful for investigating spatial organization in small brain nuclei in individual animals, which has proven difficult with multi-animal comparison approaches. We investigated cortico-subcortical projections in mice. With FluoCodes, we clearly visualized segregation of distinct cortical axons in the striatum, thalamus, pontine nuclei, brainstem, and the spinal cord. For example, in the multimodal thalamic nuclei, such as TRN, we observed topographic representation of different cortical inputs (Figure 2C). We also observed segregated inputs in ZI, another important multimodal integration hub for cortical functions (Figure 5E). Our observation may be related to how modality-specific attention and/or multimodal integration are controlled in these brain regions. In the pontine nuclei, projections from various motor and sensory areas are segregated, which may be critical for generating internal models of different sensory and motor modalities through cerebellar circuits (Figure 3E). In the Sp5 of the brainstem, we observed topographic projection from different parts of the S1, suggesting precise top-down regulation of somatosensory pathways (Figure 3F).

We also demonstrated the topographic organization of the marmoset visual system. The topographic projections from V1 neurons were compared between V2 and V5/MT. The topographic projections were also found in the two thalamic nuclei, LGN and pulvinar. Retinotopy in various visual regions has been extensively studied in macaques using classical tracer injections and electrophysiology ^58,59,70^. The combination of STPT and the FluoCode Reader, as demonstrated in this study, is an efficient strategy for studying complex 3D architecture in primates. The pulvinar is a part of the higher-order thalamus that is particularly expanded in primates. It would be interesting to investigate how inputs from V1 and higher visual areas are integrated in the pulvinar to mediate higher cognitive functions.

Previous studies have demonstrated that cortico-cortical projections are often organized in a columnar structure in the prefrontal cortex that varies considerably among individuals ^26^. Our multiplexed approach is also useful for investigating fine-scale columnar structures formed by different cortical inputs, which could be better studied through within-animal comparisons (Figure S7F). Together, our new within-animal comparison approach should be powerful for studying the organization of the higher-order brain regions in primates, contributing to our understanding of the primate-specific brain functions.

In our FluoCode system, we intentionally excluded one-XFP FluoCodes, because distinguishing between two-XFP and one-XFP FluoCodes is technically challenging (e.g., XFP1-XFP2 vs. XFP1). Chromatic aberrations and shot noise, for example, can easily produce one-XFP pixels near the two-XFP FluoCode signals. In theory, we can further increase the number of FluoCodes by using more XFPs (e.g., _8_C_2_ yields 28 combinations) and/or the combinations of three XFPs (e.g., _7_C_3_ yields 35 combinations). Additional fluorescence spectrum and lifetime variants may further expand the coding capacity of the FluoCodes in future studies ^71,72^. While we used FluoCode for projection mapping in this study, retrograde and trans-synaptic labeling could be useful for studying the organization of multisynaptic networks and loops.

FluoCode vectors may also be useful in other fields of biology. For example, they could be used for multiplexed lineage tracing in conjunction with gain-or loss-of-function experiments, particularly for morphological analyses and/or in non-traditional model organisms. Additionally, FluoCode would be a powerful tool for multiomics in future studies, integrating functional imaging *in vivo*, projectome mapping *ex vivo*, and spatial transcriptomics ^73,74^. Currently, vEM connectomics has been largely limited to small brain samples. Therefore, it would be interesting to combine the FluoCode-based whole-brain mapping approach with synaptic-scale connectomics with vEM or super-resolution microscopy in future studies ^75^.

### Limitations of the study

Identification of the FluoCode is possible only when each neuron expresses only one type of FluoCode vector. Therefore, it is important to avoid the co-expression of multiple FluoCode vectors. The injection of the AAV should be confined to a small region. It would be difficult to identify the FluoCode when multiple FluoCode AAVs infect the same neurons.

Although we can minimize the number of animals used in the study, each experiment is more time-consuming and difficult. We must prepare high-quality viruses for all of the FluoCode vectors. Injecting 21 FluoCode AAVs into one mouse typically takes four to five hours. Furthermore, performing injections in the same brain regions across multiple animals is challenging. An automated injection system would improve consistency across individuals. High-resolution imaging (seven-channel) and data analysis (21 FluoCodes) are also time-consuming processes. A high-speed imaging system is critical for applications involving mouse whole bodies and primate brains ^15,16^.

## RESOURCE AVAILABILITY

### Lead contact

Requests for additional data should be directed to and will be fulfilled on reasonable request by the Lead Contact, Takeshi Imai.

### Materials availability

The FluoCode plasmids have been deposited to Addgene (Addgene #264777-264797).

### Data and code availability

The FluoCode Reader program is available at GitHub (https://github.com/daichimori/FluoCode_Reader).

All of the image data acquired in this study will be deposited in SSBD:repository (https://ssbd.riken.jp/repository/).

Additional resources and protocols are available in SeeDB Resources (https://sites.google.com/site/seedbresources/FluoCode).

## ACKNOWLEDGEMENT

We thank M. Nishihara, T. Ohmine, E. Nozoe, S. Hamatake, D. Jeon, S. Kuriyama, F. Kamizono and M. Tsuruta for technical assistance; R. Campbell and M.Z. Lin for Addgene plasmids; H. Hashimoto, K. Seiriki, and A. Kasai for their help in building the FASTd system. We also appreciate the technical assistance from the Research Support Center, Research Center for Human Disease Modeling, Kyushu University Graduate School of Medical Sciences, which is partially supported by the Mitsuaki Shiraishi Fund for Basic Medical Research. This work was supported by Brain/MINDS (JP20dm0207055 to T.I., and JP26wm0625225 to I.I.), iBrain/MINDS (JP23wm0525012 to T.I.), and Brain/MINDS 2.0 (JP25wm0625128 to T.I. and JP24wm0625218 to A.W.), the Program for Technological Innovation of Regenerative Medicine Grant (JP26bm1123049 to I.I.), Moonshot Research & Development Program (JP22zf0127007 to I.I.), and SCARDA Kyoto University Immunomonitoring Center (KIC) (JP223fa627009 to I.I.) from Japan Agency for Medical Research and Development (AMED), CREST program of Japan Science and Technology Agency (JST) (JPMJCR2021 to T.I., and JPMJCR1921 to I.I.), the Grants-in-Aids from the Japan Society for the Promotion of Science (JSPS), KAKENHI (JP21K19355, JP21H00205, JP21H05696, JP24H02308, and JP24H02312 to T.I.; JP24K02132 to S.F.; JP24K09678 to A.W.), the SPRING Grant of JST (JPMJSP2136 to D.M.). D.M. was a predoctoral research fellow (DC1) of JSPS. B.S. was a MEXT scholar for international students.

## AUTHOR CONTRIBUTIONS

T.I. conceived the project. D.M. designed and performed most of the mouse experiments, developed the analysis pipeline, and analyzed the data. A.W. performed the marmoset experiments and the serial two-photon tomography. I.I. performed FASTd imaging. S.F., Y.I., and H.T. evaluated fluorescent proteins in HEK293T cells. B.S. helped with the construction of the data analysis pipeline. M.S. assisted with AAV production and evaluations of FluoCode vectors. H.S. developed the marmoset analysis pipeline. J.H., H.Oshiro, and H.Okano obtained *in vivo* MRI data for the marmoset. T.I. supervised the project. D.M. and T.I. wrote the manuscript with input from all the authors.

## METHODS

### Plasmids

Fluorescent proteins were initially selected based on the information in FPbase (https://www.fpbase.org/). Candidate fluorescent proteins were further evaluated based on their brightness and uniform distribution in neurons. Plasmids encoding mAmetrine1.2 (Addgene #42171) and Crimson (Addgene #190041) were obtained from Addgene. Plasmids encoding mTagBFP2 (Addgene #193339), mNeonGreen (Addgene #193341), YPet (Addgene #193342), mRuby3 (Addgene #193343), tdKatushka2 (Addgene #193344), and mTurquoise2 (Addgene #104110) were previously generated ^31^. For the construction of the FluoCode vectors, each fluorescent protein gene was PCR-amplified and subcloned into the pAAV-TRE vector. FluoCode vectors (pAAV-TRE-XFP1-P2A-XFP2-WPRE and pAAV-TRE-XFP1-EVlinker-XFP2-WPRE) generated in this study have been deposited to Addgene (Addgene #264777-264797). pmAmetrine1.2 (Addgene plasmid #42171; RRID:Addgene_42171) was a gift from Robert Campbell ^76^, and pC3.1-caggs-Crimson-CAAX (Addgene plasmid #190041; RRID:Addgene_190041) was a gift from Michael Lin ^36^.

### Mice

All mouse experiments were reviewed and approved by the Institutional Animal Care and Use Committees of Kyushu University (#A21-117 and #A23-073). C57BL/6N mice (Japan SLC, RRID: MGI: 5658686) were used for AAV experiments (8-13 week-old males and females).

### Marmosets

All marmoset experiments were reviewed and approved by the Institutional Animal Care and Use Committee of RIKEN (W2024-2-027). A four-year-old female (Figure S7F) and a five-year-old female (Figures 6, 7) marmosets were used in this study.

### Fluorescence spectrometry

HEK293T cells were cultured in high-glucose DMEM (044-29765, FUJIFILM-Wako) with 10% FBS and 1% penicillin/streptomycin (FUJIFILM-Wako) under humidified conditions in 95% air and 5% CO_2_ at 37°C. For spectral measurements, pCAG-tTA2 (0.35 μg) and one of pAAV-TRE-XFP-WPRE or pAAV-TRE-XFP1-EVlinker-XFP2-WPRE (0.9 μg) vectors were transfected into 50-80% confluent HEK293T cells in 35-mm dishes using PEI-MAX (25 mM, 5 µL/dish; Polysciences, Inc.). Twenty-four hours after transfection, cells were washed twice with 1 mL of PBS and collected with 1 mL of PBS. Cell suspensions were transferred into a glass cuvette. Excitation and emission spectra were quantified using a fluorescence spectrophotometer (F2700, Hitachi). For each XFP, spectra were measured five times in 0.5 nm wavelength increments and averaged. Excitation (Ex) and emission (Em) wavelengths to determine the spectral curves (Ex, Em) were as follows (in nm): mTagBFP2 (400, 460), mTurquoise2 (430, 505), mAmetrine1.2 (400, 535), mNeonGreen (465, 540), YPet (490, 550), mRuby3 (530, 610), Crimson (550, 650), and tdKatushka2 (550, 650). To obtain reference images for unmixing, a single type of XFP was expressed in HEK293T cells cultured on a poly-D-lysine coated 35-mm glass-bottom dish (MatTek). Twenty-four hours after transfection, cells were fixed with 4% paraformaldehyde, washed twice with PBS, and treated with SeeDB2G solution.

### Evaluations of FluoCodes in HEK293T cells

For the initial evaluation of FluoCodes in HEK293T cells, pCAG-tTA2 (0.35 μg) and one of pAAV-TRE-XFP-WPRE or pAAV-TRE-XFP1-P2A-XFP2-WPRE (0.9 μg) vectors were transfected as described above. The images were obtained by Leica TCS SP8 confocal microscopy. After linear unmixing, cell ROIs were identified with Cellpose ^77^, and the mean intensity in each of the seven XFP channels was measured for each ROI. Each channel was normalized based on its maximum value across all cells, and then scaled to unit length (L2 normalization), generating color vectors. Cells expressing a single XFP were included in the channel normalization but excluded from subsequent analyses.

FluoCodes were decoded with a nearest-centroid classifier. The centroid of the color vectors for all the transfected cells was defined as a reference for each of the 21 FluoCodes. Then, decoding was performed for each of the cells based on cosine similarity to the 21 reference color vectors (Figure S1G). Decoding accuracy was defined as the fraction of cells assigned to the ground-truth FluoCode (*n* = 8786 cells in total). The same set of cells was used for reference calculation and decoding analysis. For visualization purposes, t-SNE was used to show the segregation of color vectors (scikit-learn; perplexity = 30) (Figure S1F).

### AAV production

AAV vectors (serotype DJ) were generated using the pHelper (AAVpro Helper-Free System, Takara), pAAV-DJ (Cell Biolabs), and the AAVpro 293T cell line (#632273, Takara) following the manufacturers’ instructions. AAV vectors were purified using the AAVpro Purification Kit All Serotypes (#6666, Takara). The titration of AAV was performed using the AAVpro Titration Kit (#6233, Takara) and QuantStudio3 real-time PCR system (Applied Biosystems). AAV1-Thy1S-tTA was generated as described previously ^78^.

### AAV injections in mice

To infect AAVs, C57BL/6N mice at ages 8-13 weeks were used. Mice were anesthetized with ketamine (100 mg/kg) and xylazine (10 mg/kg) to inject a single type of AAV. Mice were anesthetized with isoflurane for long-term experiments, such as injections into over ten sites. The hair over the skull was removed and the mouse was head-fixed onto a stereotaxic apparatus (Stoelting). The scalp and underlying tissue were then removed. The skull over the injection site was thinned with a dental drill. Then, the virus was slowly injected at a rate of 5.0 nL per 10 seconds for 3 minutes (total 90 nL) using the Nanoject III system and glass capillaries (#3-00-203-G/XL, Drummond). The concentration of the injected virus cocktail was 2.5 × 10^12^ gc/mL for AAV-DJ-syn-tTA2 and 5.0 × 10^12^ gc/mL for each of the AAVDJ-TRE-XFP1-EVlinker-XFP2-WPRE. Before and after the injection, the needle was kept in place for 5 minutes. The hole in the skull was covered with Kwik-Sil (World Precision Instruments). Post-surgery, the body of the mouse was kept warm to facilitate recovery. As the toxic effects of prolonged XFP expression were observed at 4 weeks post-injection, mice were sacrificed 3-4 weeks after virus injection.

The stereotaxic coordinates of all injection sites are summarized in Table S1. In Figure 1C, injection sites for 21 FluoCodes were arrayed in a grid pattern across the dorsal cortex, each separated by ≥1.2 mm to minimize the co-infection of multiple vectors. Coordinates relative to bregma (AP, ML) and to the brain surface (DV) are shown.

### AAV injections in marmosets

We acquired a structural MRI scan under anesthesia at least one week before the virus injection. Injection sites were determined based on the presumed positions of cortical areas aligned to the Brain/MINDS reference ^79^. Structural MRI scans (T1w, T2w, DWI) were obtained using a 9.4-T BioSpec 94/30 unit (Bruker Optik GmbH, Ettlingen, Germany) and a transmit and receive coil with an 86-mm inner diameter. The stereotaxic positions were aligned by using the interaural plane and the anterior limit of the cortex. Surgery for tracer injections was performed as previously described ^80^. Pressure injection was performed using a glass micropipette with an outer diameter of 25–30 μm connected to a Nanoliter 2000 injector with a Micro4 controller (World Precision Instruments). We injected 0.25 μl of tracers at one depth (0.8 mm from the surface) over five minutes. We used a mixture of two AAVs, AAV1-Thy1S-tTA (1 × 10^9^ vg/mL) and each of the FluoCode vectors (AAVDJ-TRE-XFP1-EVlinker-XFP2-WPRE, 1 × 10^9^ vg/mL). AAV1-Thy1S-tTA has been reported previously ^78^. After surgery, the marmosets were returned to their cages and euthanized four weeks later. Stereotaxic coordinates for the injections are described in Table S1.

### Sample preparation and tissue clearing with SeeDB2G

To obtain mouse brain tissue, mice were intraperitoneally injected with an overdose of pentobarbital (P0776, TCI) in PBS for deep anesthesia, followed by intracardiac perfusion with 10 mL of PBS and 20 mL of 4% paraformaldehyde (PFA) in PBS. Removed brain samples were then fixed with 4% PFA in PBS at 4°C overnight. Brains were then embedded in 4% low-melting point agarose (ThermoFisher, #16520-100). Brain samples were cut into 520-μm-thick slices with a vibratome (PRO7N, Dosaka EM), and cleared with the SeeDB2G protocol ^81,82^. Cleared brain sections were then mounted in SeeDB2G (Omnipaque 350, Sankyo) on a glass slide using a 0.1-or 0.5-mm-thick silicone rubber sheet (AS ONE, #6-9085-12 or #6-9085-14, Togawa rubber) and glass coverslips (Marienfeld, No. 1.5H, #0109030091).

### Macroscopic fluorescence imaging

After post-fixation, brains were immersed in PBS and imaged with a fluorescence macro zoom microscope (MVX10, Olympus) (Figure 1C). Blue (mTagBFP2 and mAmetrine1.2), green (mNeonGreen and YPet), and red (mRuby3) fluorescence images were acquired using U-MWU2 (excitation 330–385 nm, emission >420 nm), U-MWIB3 (excitation 460–495 nm, emission >510 nm), and U-MWIY2 (excitation 545–580 nm, emission >610 nm) filter cubes, respectively.

### Confocal microscopy

Samples were imaged with an inverted confocal microscope, TCS SP8 (Leica) with HyD detectors using a 20× objective lens (HC PL APO 20x/0.75 IMM CORR CS2). Type G glycerol immersion (ThermoFisher, Cat#15336741) was used. We used excitation lasers at 405, 488, or 552 nm, and emission signals were dispersed by a diffraction grating as described in the legends to Figure 1C. Pinhole size was set at 1 A.U.

### Spinning disc confocal microscopy

Samples were imaged with a spinning-disc confocal microscope, Dragonfly 200 (Oxford Instruments) with a sCMOS camera, Zyla 4.2, built on an inverted microscope, ECLIPSE Ti2-E (Nikon). We used excitation lasers at 405, 488, and 561 nm and images were acquired with a 20× objective lens (CFI Plan Fluor 20XC Multi-immersion) and a spinning disc unit, CF25. Type G glycerol immersion (ThermoFisher, Cat#15336741) was used. Emission signals were dispersed by a dichroic mirror, CR-DFLY-DMQD-01I or CR-DFLY-DMPN-02I, and emission filters (center/bandwidth: 442/46, 490/60, 521/38, 540/30, 594/43, and 647/63 nm).

### FASTd imaging for whole-brain imaging

We used a custom-built FAST system (FASTd) with Dragonfly spinning-disc confocal unit (Oxford Instruments) and iQ3 software (Oxford Instruments). The FASTd system was built as described previously ^40,41^. We used excitation lasers at 405, 488, and 561 nm. Images were acquired with a 10× objective lens (Nikon CFI Plan Fluor 10X W, NA 0.3), a dichroic mirror, CR-DFLY-DMPN-02S, and emission filters (center/bandwidth: 445/46, 512/38, 521/38, 535/30, 540/20, 594/43, and 647/63 nm). Fixed brain samples were embedded in oxidized agarose as described previously ^83^. After image stitching, serial binned images were generated. Block-face sectioning was performed at 30-μm intervals, and 14 images were acquired at 5-μm z steps to cover the superficial part. The MIP of the middle 10 images per section was used to generate the whole-brain data.

### Whole-brain imaging with light-sheet microscopy

Following perfusion fixation with 4% PFA in PBS, the head and spinal column were excised. An incision to the bone was made at the cervical region, and the skull and vertebrae were carefully removed to isolate the intact brain and spinal cord. During this microdissection, the tissue was continuously hydrated with PBS. The excised intact brain and spinal cord were post-fixed in 4% PFA at 4°C for 24 hours with gentle shaking. After washing three times in PBS for 1 hour each at room temperature (RT) with gentle shaking, tissue clearing was performed using the CUBIC-L/R protocol ^46^. For delipidation, the samples were immersed in 30–40 mL of CUBIC-L solution (10 v/v% Triton X-100 [Nacalai Tesque, 35501-15] and 10 v/v% N-Butyldiethanolamine [TCI, B0725] in dH_2_O) in a 50 mL tube. The samples were incubated at 37°C for 6 days on a reciprocating shaker (50–100 rpm), with the CUBIC-L solution replaced with a fresh one on day 2. Following delipidation, the samples were washed three times in PBS for 2 hours each at RT with gentle shaking. For RI matching, the samples were first immersed in a 1:1 mixture of CUBIC-R+(M) (TCI, T3741) and Milli-Q water for 5 to 10 hours at RT with shaking in a 5 mL tube. The solution was then replaced with 100% CUBIC-R+(M), and the samples were incubated overnight at RT. To achieve optimal transparency, the 100% CUBIC-R+(M) solution was exchanged three times, each with overnight incubation at RT with shaking. After RI matching, the cleared samples were embedded in 2% (w/v) agarose [Invitrogen, 16500-500] dissolved in CUBIC-R+(M). The agarose was dissolved by heating, and the solution was kept at 65°C for approximately 1 hour to allow air bubbles to leave, since residual bubbles could cause light scattering during imaging. Embedding was performed in three layers in a custom 3D-printed mold made of PETG, with each layer allowed to solidify at 4°C for approximately 20 min before the next step. First, the bottom layer of the agarose solution was poured into the mold and allowed to solidify. Then, the sample was placed on this layer, and a second layer was added to stabilize its position. Finally, a third layer was added to cover the sample completely. After solidifying at 4°C, the gel was left at RT for approximately 3 hours to fully solidify. To obtain flat surfaces for light-sheet illumination and detection, glass slides were used for two lateral walls and placed on top of the final layer. The gel block containing the sample was immersed in CUBIC-R+(M) at RT overnight before imaging.

Cleared samples were imaged with a light-sheet microscope, UltraMicroscope Blaze (Miltenyi Biotec) with a 1.1× objective (NA 0.1, WD 17 mm) for whole-sample overviews and a 4× objective (NA 0.35, WD 16 mm) for higher-resolution imaging. During imaging, samples were immersed in a mixture of mineral oil (RI = 1.467; Sigma-Aldrich, M5904) and silicone oil (RI = 1.57; HIVAC-F4, Shin-Etsu Chemical) adjusted to a final refractive index of 1.52 to match that of the cleared tissue (CUBIC-R+(M)) ^84^. The light-sheet thickness was set to 4 μm. Samples were excited with 445, 488, and 561 nm lasers, and fluorescence was detected through the following emission filters (center/bandwidth): 480/40 nm for mTurquoise2, 510/20 nm for mNeonGreen, 545/30 nm for YPet, 595/40 nm for mRuby3, and 650/30 nm for Crimson.

The 1.1× objective was used in combination with a 0.6× optical zoom. Tiled images were acquired under the Fast Tiling mode. The z-step size was set to match the light-sheet thickness: 9.5 µm for the 1.1× objective and 4 µm for the 4× objective. BigStitcher was used to stitch the image blocks ^85^.

### Serial two-photon tomography (STPT) imaging for marmoset brains

After transcardial perfusion with 4% paraformaldehyde in 0.1 M phosphate buffer (pH 7.4), the marmoset brains were removed, post-fixed at 4°C for 2−3 days, and transferred to 50 mM phosphate buffer (pH 7.4). Agarose embedding was performed as previously described in 5% agarose using a custom-made mold. Before embedding, we treated the brain with 1 mg/ml collagenase (Wako 031-17601) at 37°C for one hour and manually removed the meninges as thoroughly as possible. STPT was performed as described using TissueCyte1600FC (TissueVision) ^86^. We immersed the agarose block in 50 mM PB supplemented with 0.02% sodium azide for imaging and sectioning. We used 920 and 1,050 nm lasers for excitation, and the two-photon excited fluorescence was detected in three-color channels (Blue Ch: <500 nm, Green Ch: 500-560 nm, Red Ch: >560 nm). In our setup, the pixel spacing for *x* and *y* was 1.390 μm/pixel and 1.420 μm/pixel, respectively. Both hemispheres were sectioned concurrently in the coronal plane and in the rostral-to-caudal direction. We imaged one optical plane every 100 μm for STPT. The entire marmoset brain was captured in 336 sections for the V1 injection sample. The processing of raw images, including stitching, 3D reconstruction and registration was performed by a pipeline used in our previous studies ^87,88^. To reconstruct the flatmap stack, the original 3D image was registered to the STPT template, and the cortical region was excised and transformed into 50 layers of cortical flatmaps, as described previously. Of the 336 sections, we collected 200 brain slices, each 100 μm thick, that exhibited any fluorescence signals. The collected sections were then mounted on glass slides (Matsunami, #S9112) using a 0.1-mm-thick silicone rubber sheet (AS ONE, #6-9085-12) as a spacer. They were then coverslipped with cover glasses (Matsunami, #C040501) and sealed with ProLong Gold antifade mountant (Thermo Fisher Scientific, #P36930). We performed 6-channel imaging of these sections with spinning-disc confocal microscopy as described above.

### Image analysis

#### 1. Preprocessing of images

To extract the fluorescence signals from a single type of XFP, we used linear unmixing as described below. For successful linear unmixing, it is important to ensure the linearity of the fluorescence intensity and low background noise. After the linear unmixing, we also subtracted leakage to the mAmetrine1.2 channel due to FRET between two fluorescent proteins.

#### 2. Linearity correction

Linearity is particularly problematic for commercialized fluorescence microscopy systems with photon-counting-based detectors, as photon detection easily saturates. In the Leica TCS SP8 with HyD detectors, the linearity is preserved in a narrow range as previously reported. Therefore, linearity correction was performed before the linear unmixing as previously described ^31^. The linearity correction program is available at GitHub (https://github.com/TakeshiImaiLab/ChromaticAberrationCorrection). It should be noted that the Leica STELLARIS system is improved but still has a similar problem. Linear range is much expanded in another photon counting system, SilVIR, used in EVIDENT confocal systems. CMOS cameras and multi-alkali PMTs preserve linearity up to the upper limit but have higher background noise.

#### 3. Background subtraction

In the CMOS camera used in the Dragonfly spinning-disc confocal imaging system (Oxford Instruments), there was a substantial level of dark noise without any fluorescence signals. Therefore, the dark noise was subtracted before linear unmixing. The dark noise was estimated from the ROIs in non-sample regions.

#### 4. Unmixing of spectrally overlapping fluorescence signals

Linear unmixing was performed as described previously ^31^. The ImageJ plugin, Linear-unmixing-QDye is available at GitHub (https://github.com/daichimori/Linear-unmixing-Qdye). Reference data were acquired with HEK293T cells expressing each of the XFPs and cleared with SeeDB2G. mTagBFP2, mTurquoise2, and mAmetrine1.2 were excited at 405 nm and fluorescence at 410-468 nm (Ch1), 468-505 nm (Ch2), and 520-600 nm (Ch3) were acquired. mNeonGreen and YPet were excited at 488 nm, and fluorescence at 495-525 nm (Ch4) and 525-555 nm (Ch5) were acquired. mRuby3 and Crimson (or tdKatushka2) were excited at 552 nm and fluorescence at 575-600 nm (Ch6) and 615-755 nm (Ch7) were acquired.

In brain sample applications, references were obtained directly from injection sites of the sample. In this case, it is important to select ROIs that have only a single type of FluoCode. In ImageJ, a rectangular ROI was drawn around each injection site, and labeled and unlabeled pixels were separated by Otsu thresholding within the ROI. The fluorescence ratios across channels were then used as reference parameters for linear unmixing. In this way, we calculated intensity data, *I_X_*, for each of the XFPs.

#### 5. Subtraction of two types of autofluorescence signals using linear unmixing (for marmoset brains only)

Marmoset brain samples exhibited stronger autofluorescence than the mouse brain. We therefore employed additional preprocessing steps to subtract autofluorescence. Two distinct types of autofluorescence were observed: a diffuse neuropil-specific autofluorescence with a short-wavelength emission profile, and lipofuscin-derived autofluorescence with a broader, longer-wavelength emission profile. The diffuse neuropil-specific autofluorescence was stronger in the marmoset, possibly due to prolonged post-fixation and/or intrinsic properties in marmosets. Lipofuscin is an autofluorescent pigment that accumulates with age in the brain and is particularly abundant in primates.

To obtain FluoCode signals, neuropil-and lipofuscin-derived autofluorescence were subtracted by two additional rounds of linear unmixing. Reference data for the autofluorescence were obtained from ROIs without any XFP signals. For each ROI, lipofuscin-containing and lipofuscin-free pixels were separated by Otsu thresholding using a long-wavelength lipofuscin channel (excitation 405 nm, emission 573–616 (594/43 filter) nm). For lipofuscin-free pixels, we used the mTagBFP2 channel (excitation 405 nm, emission 442/46 nm) to obtain the reference data for neuropil autofluorescence. For lipofuscin-containing pixels, fluorescence in the lipofuscin channel (excitation 405 nm, emission 573– 616 nm (594/43 filter)) was used for lipofuscin autofluorescence; the contribution of neuropil autofluorescence was negligible. In the fluorescence imaging of the FluoCode, we obtained two additional channels, mTagBFP2 (for neuropil autofluorescence) and lipofuscin channel. Using the reference data for XFP and the two types of autofluorescence (neuropil, lipofuscin), we performed linear unmixing to separate XFP signals (*I_X_*) vs. two types of autofluorescence (Figure S7).

#### 6. Subtraction of FRET-derived signals

For the four specific FluoCodes (BN, BY, NT, and YT), we observed Förster resonance energy transfer (FRET) between a blue-excited XFP (mTagBFP2 or mTurquoise2) and a green-excited XFP (mNeonGreen or YPet), producing spurious bleed-through into the mAmetrine1.2 channel (but not to other channels).

To remove the FRET-based leakage to the mAmetrine1.2 channel, we defined the leakage level from the four FluoCode a-b (*L_a,b_*) as follows:

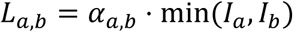

where *α_a,b_* is an optimized scaling factor for each donor–acceptor pair. The value of *α_a,b_* for each pair was estimated from the reference images (e.g., injection site). Based on mAmetrine1.2 signal, *I_A_*, and *L_a,b_* values, the corrected intensity for mAmetrine1.2, *I_A*_*, was calculated as follows:

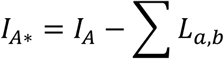

In this way, we obtained intensity data, *Ix*, for all the XFPs, without background and autofluorescence contaminations. We did not perform vector normalization of *I* for different XFP signals prior to FluoCode Reader, but *Ix* for different XFPs was typically within a 3-fold difference.

### FluoCode Reader program

Each FluoCode is characterized by the co-expression of two specific XFPs (a and b) out of seven. The FluoCode Reader classifies each pixel to one of the 21 FluoCode channels. The FluoCode Reader pipeline consists of hue extraction (Step 1), probabilistic extraction of morphological features (Step 2), and final FluoCode assignment (Step 3).

#### Step 1: Hue extraction

For each FluoCode defined by two specific channels (XFP-a and -b), we calculated hue-extracted images (*I_ab_*) based on seven XFP images, *I_X_*, as follows:

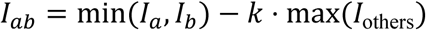

Where *I_a_, I_b_*, and *I_others_* indicate the signal intensity for channel a, b, and all the others, and min (*I_a_, I_b_*) is the minimum intensity (pixel-wise) between the two target channels and max (*I_others_*) is the maximum intensity (pixel-wise) of the non-target channels. *k* is a leakage parameter and was set to minimize the residual crosstalk across channels. The parameter *k* was heuristically set to 0.2 in our analyses, because the leakage across channels did not exceed 10% after unmixing in our data (Figure 1B). Negative values were clipped to zero. In this way, we performed hue extraction for all the 21 FluoCodes.

#### Step 2: Probabilistic extraction of target signals

For each FluoCode defined by XFP-a and -b, a random forest classifier trained in ilastik (https://www.ilastik.org/)^54^ was applied to *I_ab_* (from Step 1) to generate a probability map, *P_ab_* representing the probability that the pixel contains specific morphological features (i.e., neurons and neurites). This procedure was performed for all 21 FluoCodes. Details of the classifier training are described in the following section (Training the random forest classifier for the FluoCode Reader).

It should be noted that *P_ab_* may pick up target features present in only one of the XFP channels when the background level is higher for the other XFP channel. To ensure that we pick up specific morphological features present in both of the XFP channels, we also generated probability maps, *P_X_*, based on all the seven XFP images, *I_X_*, and the training data using ilastik. The corrected probability map, *P_ab*_*, was then calculated as follows:

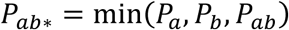

*P_ab*_* will only pick up specific morphological features present in both of the XFP images, *I_a_* and *I_b_*.

#### Step 3: FluoCode assignment

In this way, we calculated a probability map, *P_ab*_*, for all 21 FluoCodes. For each pixel, we then identified the FluoCode with the maximum *P_ab*_*, based on argmax. Here, we considered *P_ab*_* below the heuristic threshold value of 0.2 as background. In this way, we generated decoded FluoCode images, in which the decoded FluoCode was represented by numbers (No FluoCode = 0; FluoCode-1 = 1; FluoCode-2 = 2; …).

The FluoCode Reader is implemented as a Fiji/ImageJ plugin, available at our GitHub repository (see Data and code availability).

### Training the random forest classifier for the FluoCode Reader

The ilastik classifier was trained using images from all seven XFP channels. For the training with manual annotation, 1-2 representative *x-y* images per XFP channel were selected. These images were chosen to contain a wide range of morphological features, such as neuronal somata (injection sites), dendrites, thick axonal bundles, and fine axonal fibers, to detect diverse morphological features. A single classifier was used for all the FluoCode images of the same experiment. We confirmed successful segmentation across XFP channels and FluoCodes. The ilastik project files (.ilp), including the trained classifiers, hyperparameters (e.g., number of trees and selected features), training images, and manual annotations, used in our study are archived on GitHub (see Data and code availability). We used ilastik for all the segmentations; however, our FluoCode Reader is also compatible with LabKit, another Fiji/ImageJ plugin using the same random forest-based segmentation algorithm.

### Registration of the brain regions

Whole-brain image registration to the Allen Mouse Brain Common Coordinate Framework (CCFv3, https://alleninstitute.github.io/abc_atlas_access/descriptions/Allen-CCF-2020.html)^7^ was performed in two steps. First, we performed a coarse registration using a Fiji/ImageJ plugin, Elastix (https://imagej.net/plugins/elastix). We then performed manual refinement using a Fiji/ImageJ plugin, BigWarp (https://imagej.net/plugins/bigwarp). We deformed the reference coordinate, rather than the raw image data, for the registration. For the two-dimensional images of brain sections, the corresponding coronal planes were first selected from CCFv3. The selected sections of CCFv3 were then manually deformed and registered to the fluorescence image data using BigWarp.

### Quantification of projection density

Projection density for each brain region was quantified from the decoded FluoCode images. After registration of CCFv3 to the decoded FluoCode images, we counted the number of pixels classified as the target FluoCode within each region. Projection density for a FluoCode a-b (*D_ab_*) was defined as the fraction of the FluoCode a-b-positive pixels relative to the total number of pixels within the brain region in CCFv3:

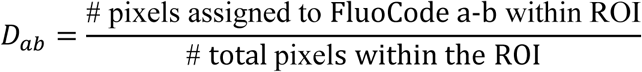

Pixel counting per region was performed using a custom ImageJ plugin. The plugin source code is available at our GitHub repository (see Data and code availability).

In Figure 5, CR was excluded from the quantitative analyses because its expression level was quite low in this animal.

### Comparison with the Allen Mouse Brain Connectivity Atlas

For each of the 14 FluoCode injection sites, a corresponding experiment was selected from the Allen Mouse Brain Connectivity Atlas based on the similarity in coordinate information. We only considered wild-type experiments. The selected experiment IDs are listed in Table S2.

Projection data were retrieved from the Allen Institute RMA API (http://api.brain-map.org/api/v2/data/query.json) using the ProjectionStructureUnionize. Structure IDs were resolved from acronyms within graph_id = 1. The projection metric was projection density, and records with hemisphere_id = 3 (both hemispheres) were used, because pixels from both hemispheres were counted in our FluoCode dataset. Brain regions were defined and grouped as in our FluoCode dataset, so that each dataset was represented as a matrix of 34 thalamic regions × 14 injections.

Cross-correlation matrix (Spearman’s rank correlation) was used to evaluate the similarity between Allen and FluoCode datasets. This result was also compared with FluoCode × FluoCode and Allen × Allen comparisons.

### 3D segmentation of FASTd, light-sheet, and STPT images

Because the TissueCyte STPT acquires only three fluorescence channels, we could not identify FluoCodes directly from the 3D STPT images. Instead, we used FluoCode Reader to identify FluoCodes in brain sections. We then manually annotated the FluoCodes in the STPT volume data using Imaris 10.1 (Oxford Instruments). A region of interest (ROI) was manually defined for the target signals in the STPT volume. The machine learning–based “Surface Generation” module of Imaris was used to surface render the neuronal structures in 3D. Similarly, we used the Surface Generation module of Imaris 10.1 to manually generate surface rendering for FASTd and light-sheet images.

### Quantification of axonal segregation in marmoset V2 and V5/MT

To compare the segregation of axonal projections in V2 and V5/MT, we analyzed pairs of FluoCodes whose terminal fields were adjacent to each other in the same sections in both areas. We used the five FluoCode pairs (RA–CY, CN–CY, NT–YN, YN–RA, and YN–CY) which met this criterion (Table S4).

For each pair and each area, an ROI with a 300-µm width was drawn manually along the cortical surface using Fiji/ImageJ. Then the intensity profile of each FluoCode along the cortical surface was obtained with Plot Profile. Each profile was smoothed twice with a three-point moving average, and FluoCode signals below 1,000 were excluded as background. Each remaining profile was then normalized to its maximum value (Figure 6E).

*Overlap coefficient.* The normalized intensity profiles of FluoCode-1 and -2 at position *x* were defined as *f*_1_(*x*) and *f*_2_(*x*), respectively. Then, the overlap coefficient (*OC*) was defined as follows:

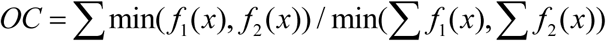

The OC value is 1 when one profile lies entirely within the other and 0 when the two profiles do not overlap at all.

*Separation index.* For FluoCode-1 and -2, the median positions (*M_1_* and *M_2_*) and interquartile ranges (*IQR_1_* and *IQR_2_*) were calculated. The separation index (*SI*) was defined as follows:

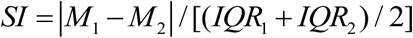

A larger *SI* value indicates that the two projections are further apart relative to their spatial spreads.

### Statistical analysis

*OC* and *SI* were compared between V2 and V5/MT using the two-sided paired *t*-test (*n* = 5 FluoCode pairs from one marmoset). *P* < 0.05 was considered statistically significant.

For the comparison with the Allen Mouse Brain Connectivity Atlas, Spearman’s rank correlation coefficients were calculated across *n* = 34 thalamic regions for each of the FluoCode–Allen pairs. Significance was evaluated using the *t* approximation with the Holm method.

Statistical analyses were performed with Python and SciPy.

## SUPPLEMENTAL INFORMATION

**Figure S1.**
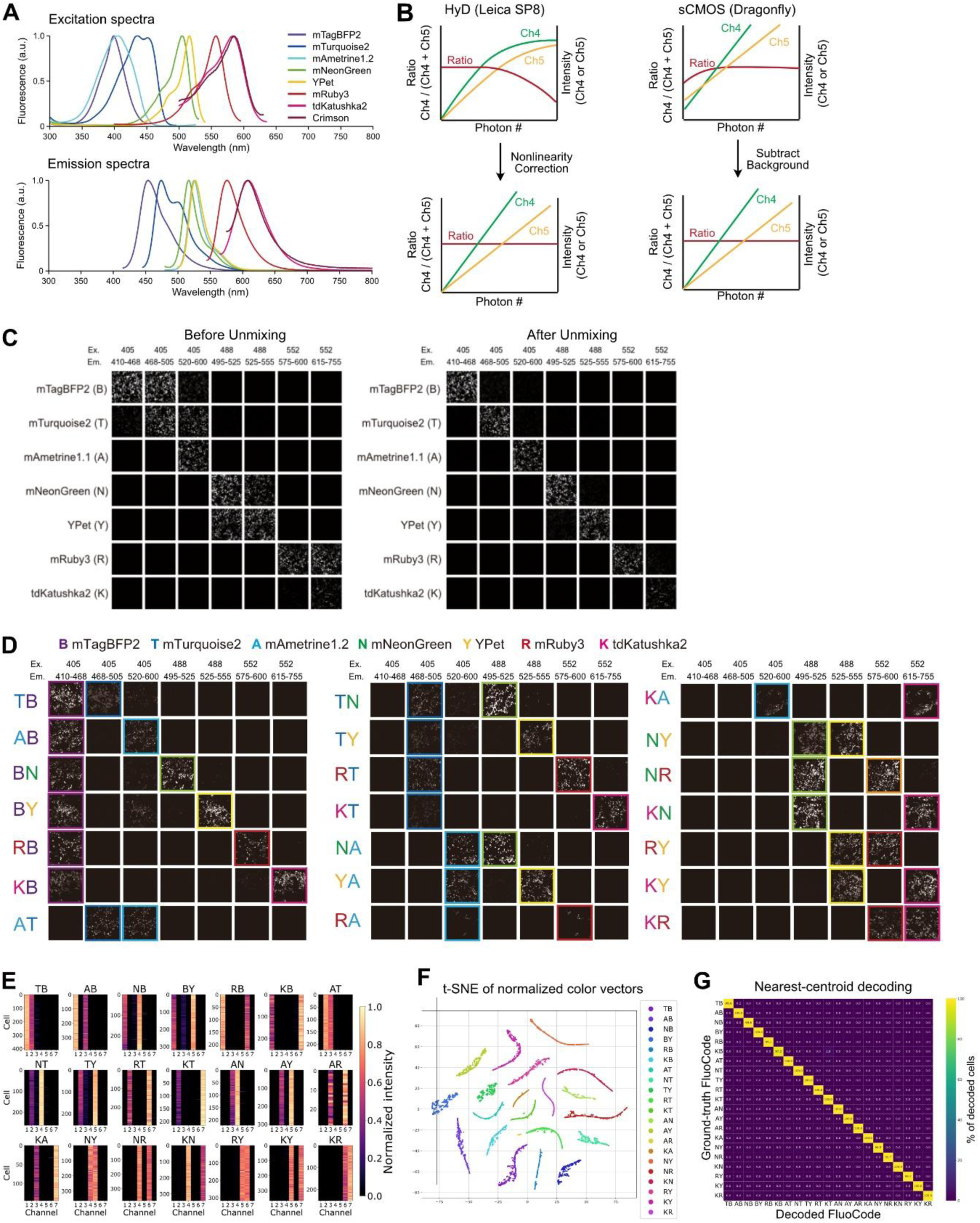
Evaluation of the 21 FluoCodes in HEK293 cells, related to Figure 1. (A) Excitation (top) and emission (bottom) spectra of the seven XFPs selected for the FluoCode vectors. We used lasers at 405 nm (mTagBFP2, mTurquoise2, and mAmetrine1.2), 488 nm (mNeonGreen and Ypet), and 552/561 nm (mRuby3 and Crimson/tdKatushka2) for excitation. Their emission peaks were sufficiently separated to allow for linear unmixing across seven channels. (B) Schematic of the image data preprocessing required for accurate linear unmixing. The fluorescence ratios between channels (e.g., Ch4/(Ch4 + Ch5)) must be constant for successful linear unmixing. Left: With the photon-counting HyD detectors of the Leica TCS SP8, signal intensity saturated at high photon numbers. We used a linearity correction program as reported previously ^31^. Right: With the sCMOS camera of the Dragonfly spinning-disc confocal system, dark noise was present. We subtracted the background signals to ensure linearity. (C) Linear unmixing separates the signals of seven XFPs expressed in HEK293 cells. Each XFP was imaged under seven different excitation/emission conditions (Ex/Em, nm). The left panel shows the fluorescence signals before unmixing, demonstrating some spectral overlap across the XFPs. The right panel shows the images after unmixing, demonstrating clear separation of the XFP signals. (D) Validation of the 21 FluoCode vectors in HEK293 cells following linear unmixing. Each FluoCode was imaged under seven different excitation/emission conditions. FluoCodes are labeled with two-letter abbreviations (e.g., TB indicates co-expression of mTurquoise2 and mTagBFP2). Colored boxes highlight the two channels with the expected XFP signals. All 21 FluoCodes were identified in an all-or-none fashion after linear unmixing. Note that we used XFP1-P2A-XFP2 constructs in our initial evaluation with HEK293 cells. For better localization in *in vivo* studies, we then switched to XFP1-EV linker-XFP2 constructs (Figure 1). (E) Color vectors of HEK293 cells expressing each FluoCode. Regions of interest (ROIs) for cells were identified with Cellpose, and the XFP signals for each ROI (after unmixing) were vector normalized. Channels 1–7 correspond to the excitation/emission combinations shown in (C) and (D). (F) t-SNE plot of the vector-normalized color vectors for HEK293 cells expressing each FluoCode. Cells expressing the same FluoCode formed a discrete cluster. (G) Confusion matrix of a linear decoder for FluoCodes. Vector-normalized cell signals were used for the nearest centroid linear decoder ^31^. The ground truth refers to the transfected FluoCode vectors. The values indicate the percentage of cells assigned to each FluoCode (decoding accuracy = 99.7%).

**Figure S2.**
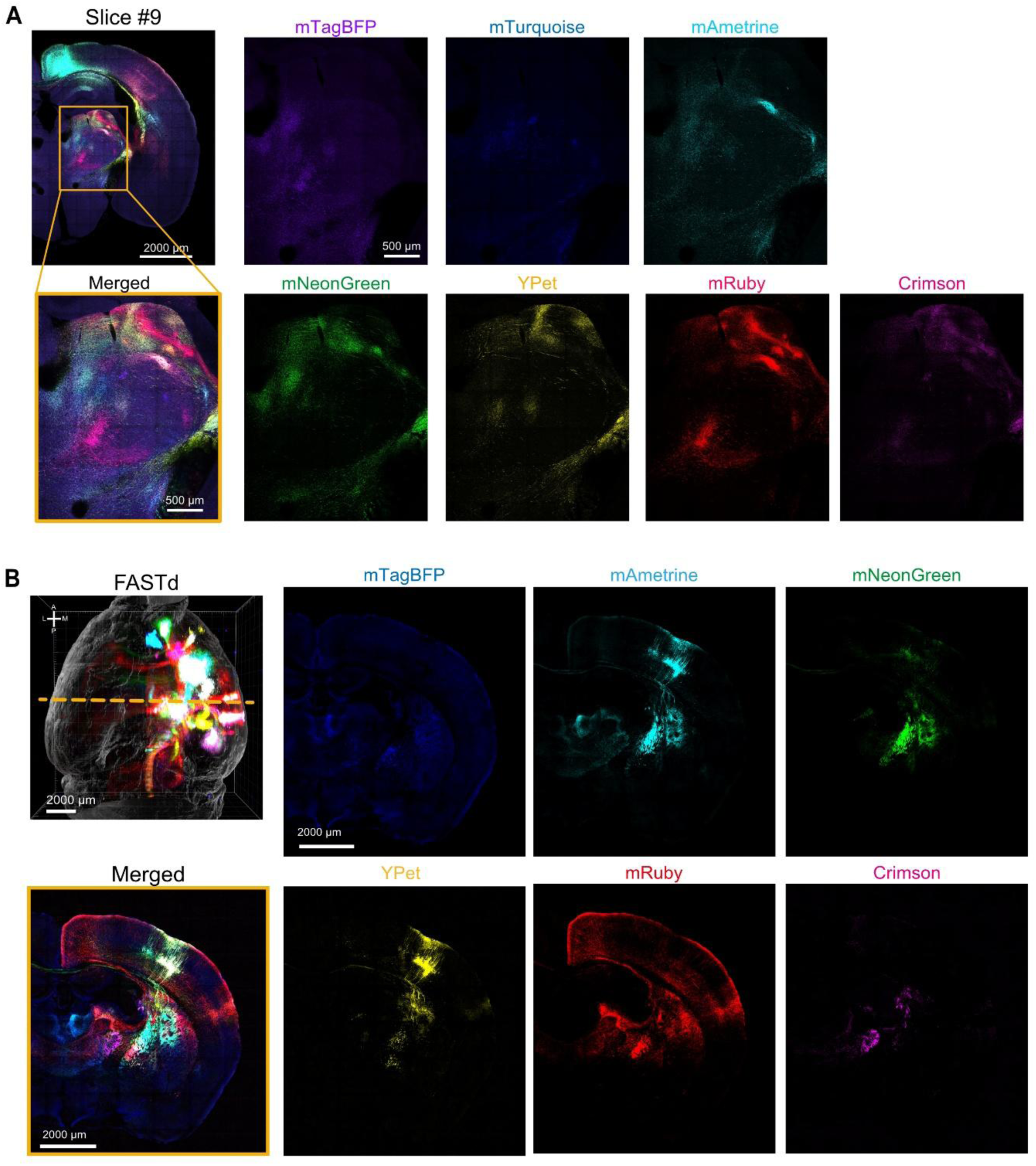
Single-channel images of FluoCode-labeled brain samples, related to Figures 1 and 2. (A) Individual XFP channels of the coronal section shown in Figure 1E (slice #9, left). Scale bars, 2,000 µm (upper left) and 500 µm (bottom left and right). (B) Individual XFP channels of a FASTd image. The top left shows the 3D rendering of the entire brain shown in Figure 2A. The dotted line indicates the location of the coronal section. Scale bars, 2,000 µm.

**Figure S3.**
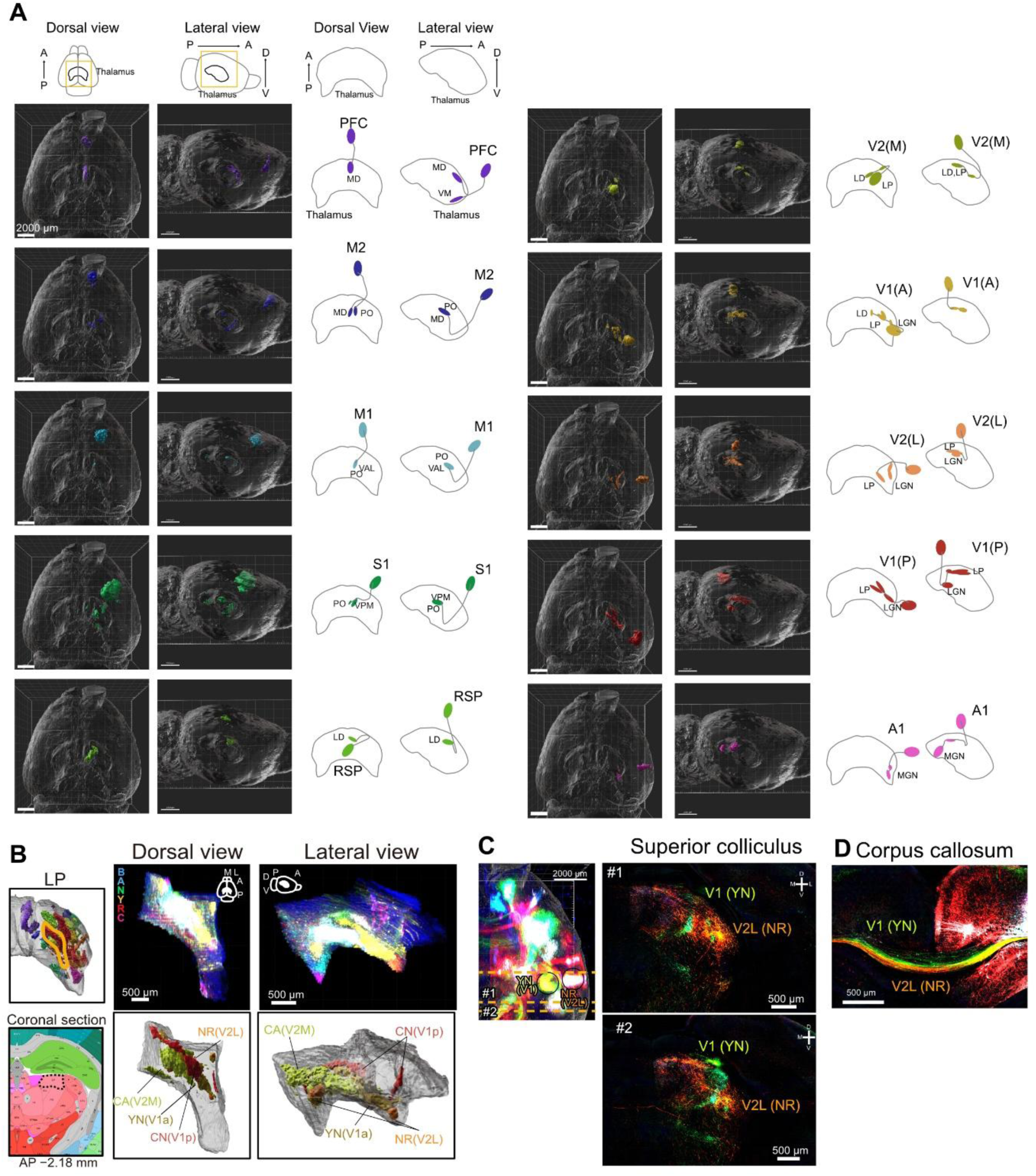
Topographic organization of corticofugal projections revealed by FASTd, related to Figure 2. (A) Corticothalamic projections from individual cortical areas. The 3D reconstruction of labeled axons from each of the ten representative injection sites (PFC, M2, M1, S1, RSP, V2M, V1(A), V2L, V1(P), and A1) is shown in horizontal (left) and sagittal (middle) views. The schematics summarize the thalamic nuclei targeted by each cortical area. A, anterior; P, posterior; D, dorsal; V, ventral. Scale bars, 2,000 µm. (B) Topographic organization within the lateral posterior nucleus (LP; left) and the posterior medial nucleus (POm; right). Their positions are indicated in the thalamic reconstruction (orange boxes) and in the corresponding coronal section of the Allen CCFv3 atlas (black dotted lines; AP, −2.18 mm from bregma). Dorsal and lateral views of fluorescence (top) and segmentation (bottom) are shown on the right. The FluoCode signals for the visual areas found in the POm are most likely derived from a spillover of the injections into the neighboring S1. Scale bars, 500 µm. (C) Axonal projection to the superior colliculus. Coronal sections show that axons from V1 and V2L terminate in distinct domains within the superior colliculus. The left image shows a dorsal view of the 3D rendering showing the positions of the coronal planes in (C) (#1 and #2) and (D). Circles indicate the injection sites. Scale bar, 500 µm. (D) Topographic organization of callosal axons. Axons from V1 (YN) and V2L (NR) travel through distinct parts of the corpus callosum. Scale bar, 500 µm.

**Figure S4.**
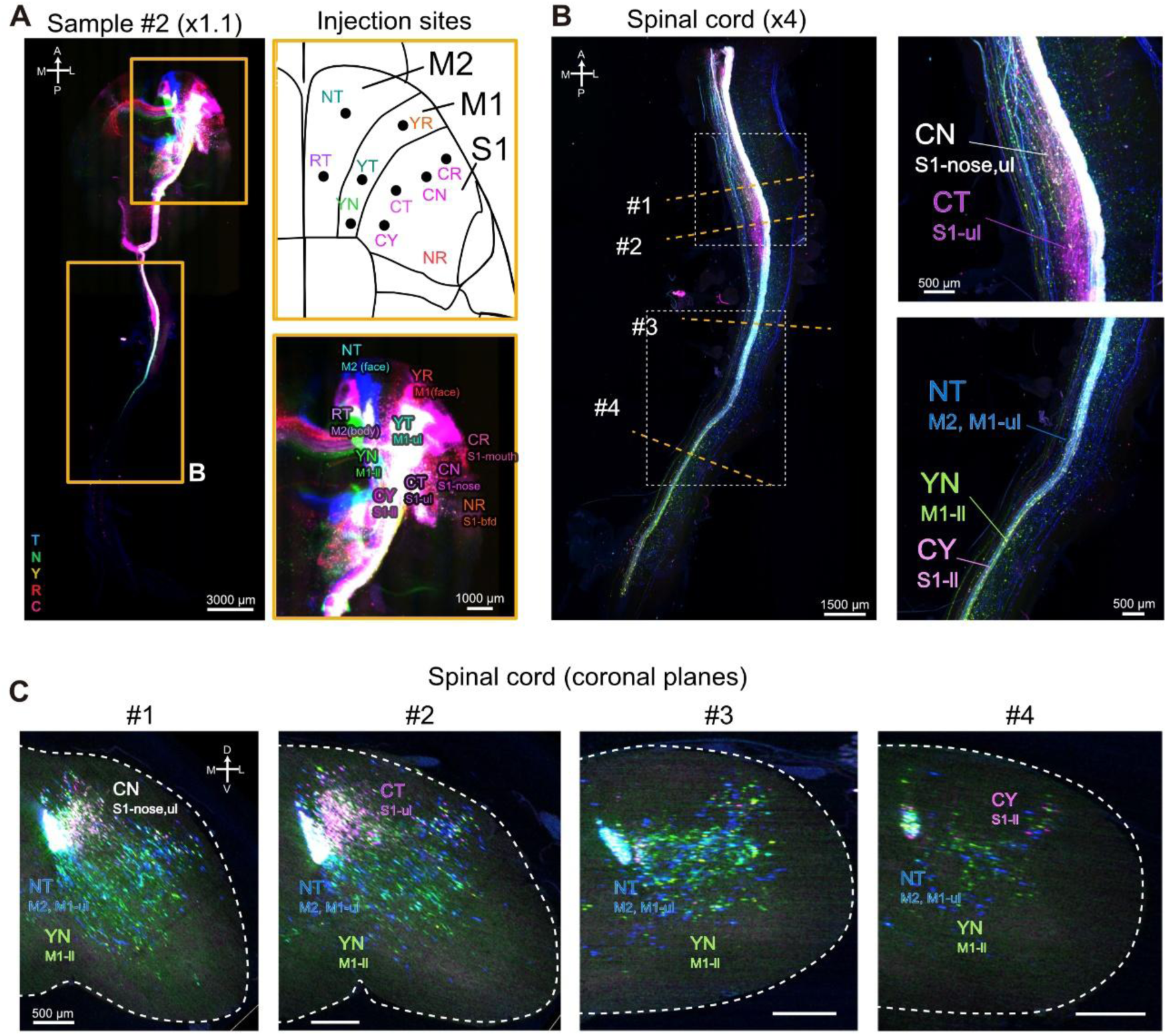
Topographic organization of corticospinal axons, related to Figure 3. (A) A dorsal view of another sample (Sample #2), imaged using light-sheet microscopy with a 1.1× objective. The injection sites for 10 FluoCode vectors are shown on the right. We targeted motor and somatosensory areas (NT, RT, YT, YN, YR, CT, CN, CR, CY, and NR). ul, upper limb; ll, lower limb; bfd, barrel field; A, anterior; P, posterior; M, medial; L, lateral. Scale bars, 3,000 µm (left) and 1,000 µm (lower right). (B) Dorsal views of the spinal cord imaged with a 4× objective. Orange dotted lines (#1–#4) indicate the positions of the coronal planes shown in (C). Magnified views of the boxed regions are shown on the right. Axons from the S1 nose/upper limb (CN) and S1 upper limb (CT) were found in the rostral part. Axons from the M2/M1 upper limb (NT), M1 lower limb (YN), and S1 lower limb (CY) were found in the caudal region. Scale bars, 1,500 µm (left) and 500 µm (right). (C) Coronal planes of the spinal cord at the levels indicated in (B), from rostral (#1) to caudal (#4). Dotted lines mark the contralateral hemicord. S1 axons (CN, CT, and CY) mainly terminated in the dorsal horn. M1/M2 axons (NT and YN) were distributed in the intermediate zone, the ventral horn, and the medial dorsal horn. D, dorsal; V, ventral; M, medial; L, lateral. Scale bars, 500 µm.

**Figure S5.**
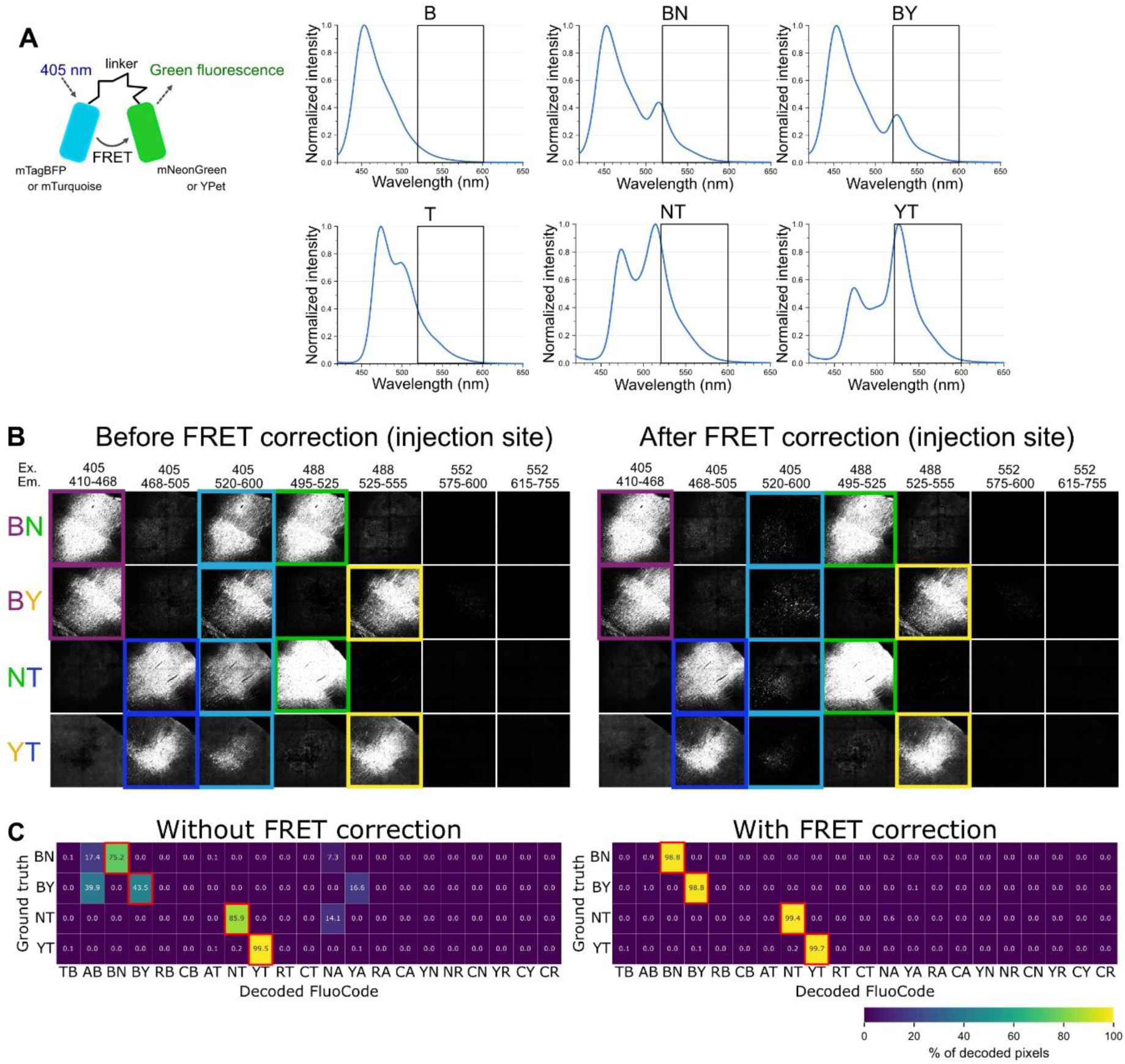
FRET correction for the FluoCode Reader, related to Figure 4. (A) FRET between two XFPs in the FluoCode. FRET was observed between a blue donor (mTagBFP2 or mTurquoise2) and a green/yellow acceptor (mNeonGreen or YPet), which were connected by an EV linker. Excitation of the donor at 405 nm produces emission from the acceptor. The emission spectra of FluoCodes under 405-nm excitation are shown. The spectra were obtained from HEK293 cells expressing the donor alone (B and T) or donor– acceptor pairs (FluoCodes; BN, BY, NT, and YT). Black boxes indicate the mAmetrine1.2 channel detection window (Em 520–600 nm). (B) FRET correction for BN, BY, NT, and YT. Images of the injection sites (shown in Figure 1B) are shown before (left) and after (right) FRET correction. Colored boxes indicate the detected signals after unmixing. In addition to the signals in the expected channels, FRET-derived signals were found in the mAmetrine channel (Ex 405/Em 520–600 nm). The FRET-derived signals were removed after correction (See Methods for details). (C) Confusion matrices of the FluoCode Reader before (left) and after (right) FRET correction (BN, BY, NT, and YT). Red boxes indicate the ground-truth FluoCodes. The values indicate the percentage of pixels assigned to each FluoCode.

**Figure S6.**
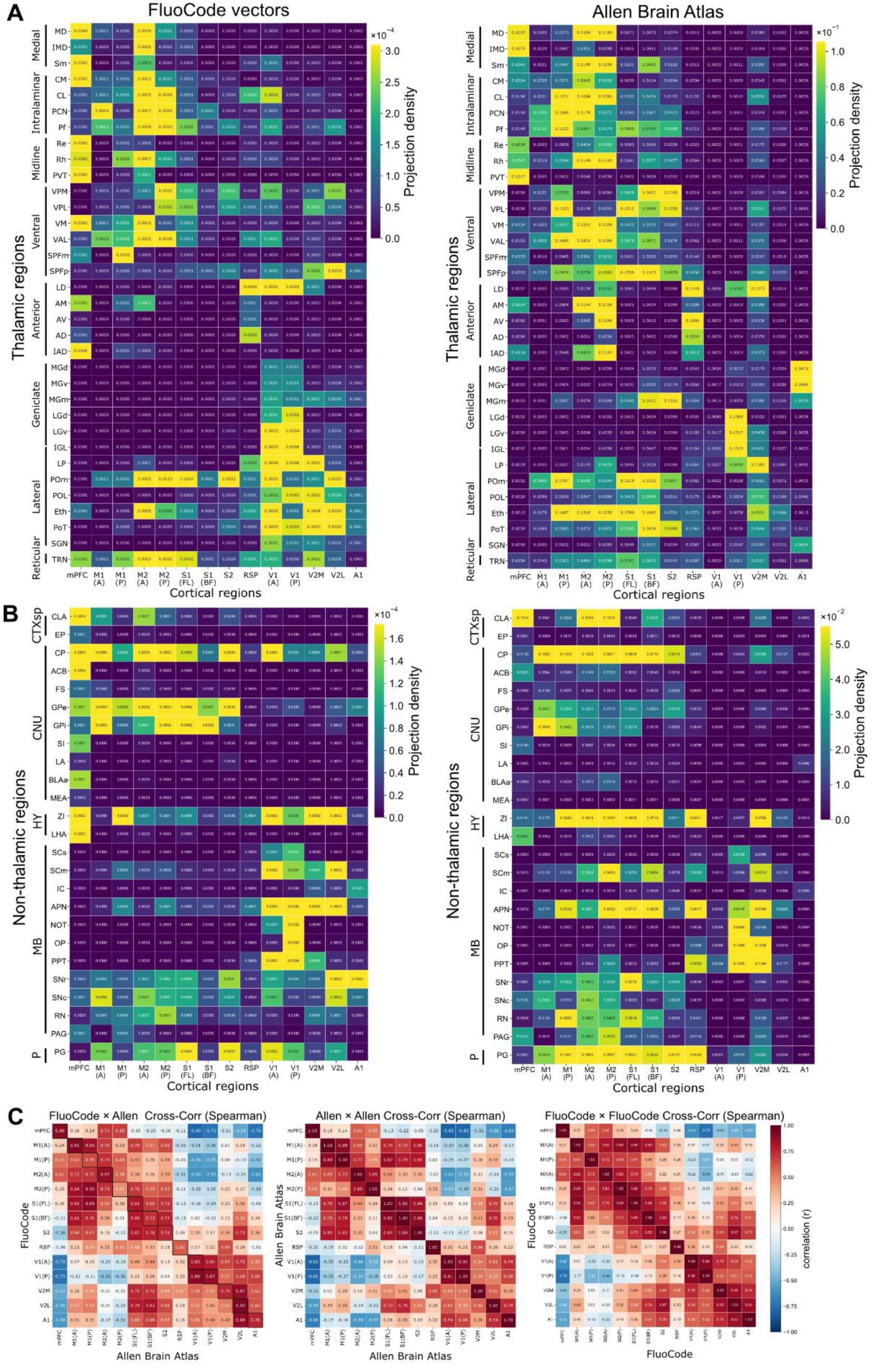
Comparison with the Allen Mouse Brain Connectivity Atlas, related to Figure 5. (A) Projection density in thalamic nuclei for 14 cortical injection sites, obtained using FluoCode vectors (left) and the Allen Mouse Brain Connectivity Atlas (right). For each FluoCode injection site, the closest Allen experiment was selected (experiment IDs are listed in Table S2). The thalamic nuclei are grouped by subdivisions (medial, intralaminar, midline, ventral, anterior, geniculate, lateral, and reticular). (B) Projection density in non-thalamic subcortical regions. Target regions are grouped by major subdivisions: cortical subplate (CTXsp), cerebral nuclei (CNU), hypothalamus (HY), midbrain (MB), and pons (P). (C) Spearman’s rank correlation of projection profiles across thalamic regions. Comparisons were made for FluoCode × Allen data (left), Allen × Allen, and FluoCode × FluoCode. Black boxes in the left matrix indicate matched pairs. mPFC, medial prefrontal cortex; M1(A)/M1(P), anterior/posterior part of M1; M2(A)/M2(P), anterior/posterior part of M2; S1(FL), S1 forelimb region; S1(BF), S1 barrel field; RSP, retrosplenial agranular cortex; V1(A)/V1(P), anterior/posterior part of V1. Abbreviations of brain regions are summarized in Table S3.

**Figure S7.**
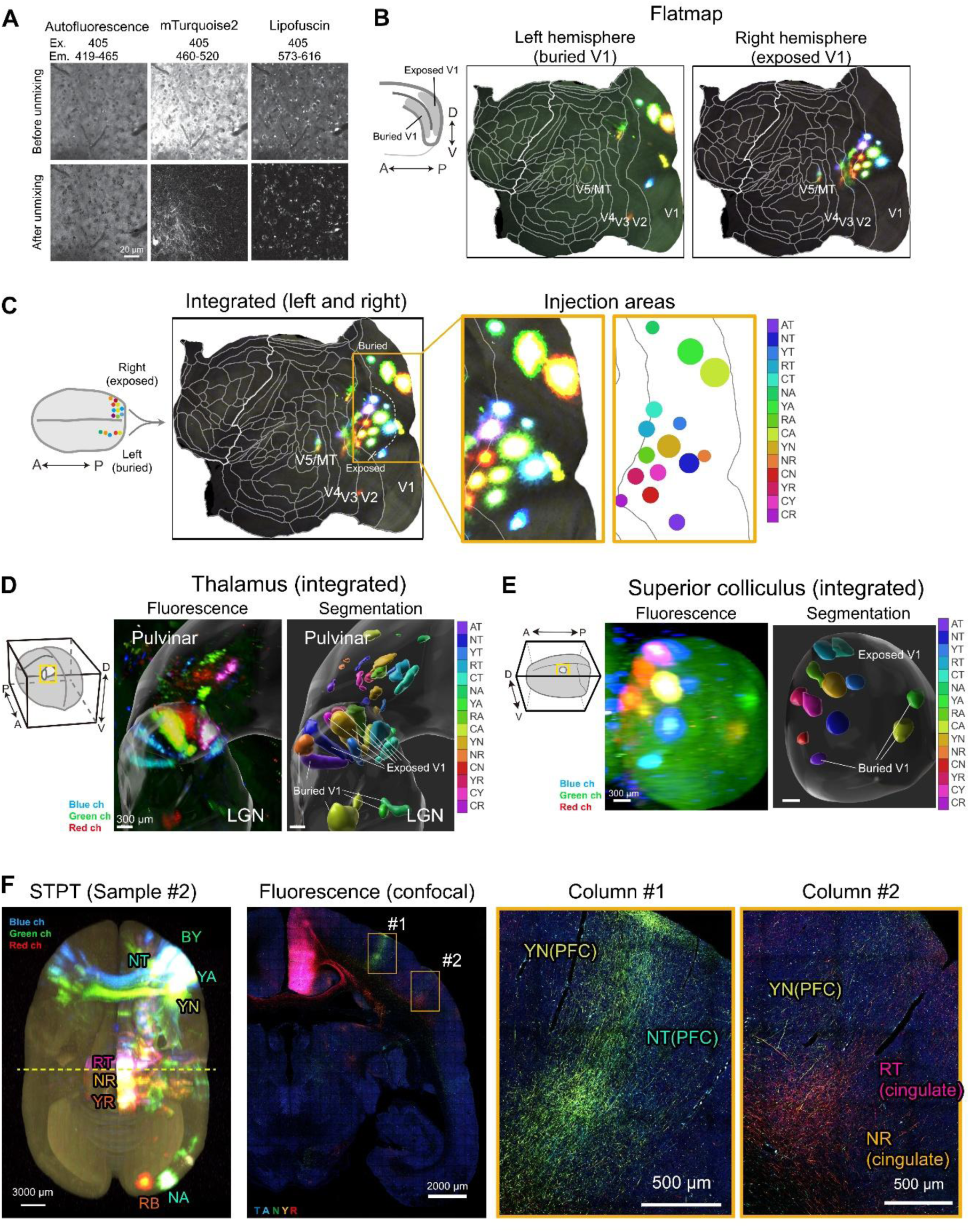
Projectome analysis of the marmoset brain, related to Figures 6 and 7. (A) Removal of tissue autofluorescence and lipofuscin signals by linear unmixing. A marmoset cortical section without any fluorescent proteins was imaged by confocal microscopy using 405 nm excitation and three emission bands: 419–465 nm (tissue autofluorescence), 460–520 nm (mTurquoise2), and 573–616 nm (lipofuscin). Before unmixing (top), the mTurquoise2 channel contained considerable tissue autofluorescence and lipofuscin signals. After unmixing, these signals were fully removed from the mTurquoise2 channel (bottom). Scale bar, 20 µm. (B) Cortical flatmaps ^26^ reconstructed from serial two-photon tomography (STPT) data of sample #1 (Figures 6 and 7), showing FluoCode injection sites in V1. Multiple FluoCodes were injected into the buried V1 regions of the left hemisphere (left) and into the exposed V1 regions of the right hemisphere (right). The schematics on the left show the positions of the buried and exposed regions of V1. White lines indicate cortical areal boundaries, and V1, V2, V3, V4, and V5/MT are labeled. D, dorsal; V, ventral; A, anterior; P, posterior. (C) Cortical flatmap of the integrated FluoCode signals from both hemispheres of sample #1 (left). The data from the left hemisphere (buried V1) were mirrored onto the right hemisphere (exposed V1), so that all 15 injection sites are represented on a single flatmap. The dotted line delineates the buried and exposed V1 regions. Magnified views of the injection areas are shown as a fluorescence image (middle) and a color-coded schematic (right). White lines delineate areal boundaries, and V1, V2, V3, V4, and V5/MT are labeled. A, anterior; P, posterior. (D) 3D rendering of V1 projections to the thalamus (LGN and pulvinar) based on the STPT data. The orientation and the location of the displayed region are shown schematically on the left (yellow box). The segmentation shows the terminal fields that originate from buried and exposed V1 injections. Scale bars, 300 µm. (E) 3D rendering of V1 projections to the superior colliculus (SC), based on STPT data. Scale bars, 300 µm. (F) Mapping of projections from the prefrontal cortex (PFC), the cingulate cortex, and V1 in Sample #2 using FluoCode vectors. Injection sites: PFC (BY, NT, YA, and YN), the cingulate cortex (RT, NR, and YR), and V1 (RB and NA). A dorsal view of the whole brain imaged with STPT is shown on the left. The yellow dotted line indicates the position of the coronal section shown in the middle. This section was imaged with six-channel spinning-disc confocal microscopy. Axon terminals often demonstrated a columnar architecture ^26^, and axons labeled with different FluoCodes project to the same column. FluoCodes from two PFC sites (YN and NT) in column #1 and from PFC (YN) and the cingulate cortex (RT and NR) in column #2. Scale bars, 3,000 µm (left and middle) and 1,000 µm (right).

### Supplementary Table

**Table S1.** Stereotaxic coordinates and FluoCode assignments for AAV injections in the mouse and marmoset cortex. Injections of 21 FluoCodes were arrayed in a grid pattern across the dorsal cortex, separated by ≥1.2 mm, and used for whole-brain imaging of coronal sections (Figures 1C–E); cortex-wide injections were used for the FASTd experiment (Figure 2) and quantitative connectivity mapping (Figures 5, S6); motor and somatosensory injections were used for imaging with light-sheet microscopy (Figure 3); marmoset V1 injections were used for serial two-photon tomography of sample #1 (Fig. 6 and 7). Anteroposterior (AP) and mediolateral (ML) coordinates are given relative to bregma, and depth is given from the brain surface at each injection site. All mouse injections were performed unilaterally in the right hemisphere; in the marmoset, the exposed V1 was injected in the right hemisphere and the buried V1 within the calcarine sulcus in the left hemisphere. ML values are shown as absolute distances from the midline. (A) and (P) denote anterior and posterior injection sites within the same area. Cortical areas: mPFC (PrL), prelimbic region of the medial prefrontal cortex; M1, primary motor cortex; M2, secondary motor cortex; S1, primary somatosensory cortex; S2, secondary somatosensory cortex; RSP, retrosplenial agranular cortex; V1, primary visual cortex; V2M, secondary visual cortex, medial part; V2L, secondary visual cortex, lateral part; A1, primary auditory cortex; AuD, secondary auditory cortex, dorsal area; TeA, temporal association cortex; PtA, parietal association cortex; FL, forelimb; BF and bfd, barrel field; ul, upper limb; ll, lower limb. A slash (e.g., V2M/PtA) indicates a site at the border between two areas.

| FluoCode | Cortical area | AP (mm) | ML (mm) | Depth (mm) |
|---|---|---|---|---|
| *Injections of 21 FluoCodes (Figure1)* |  |  |  |  |
| TB | M2(A) | +2.4 | 0.7 | 0.6 |
| AB | M1(A) | +2.4 | 1.8 | 0.6 |
| BN | M2(A) | +1.2 | 0.7 | 0.6 |
| BY | M1(P) | +1.2 | 1.8 | 0.6 |
| RB | S1-mouth | +1.2 | 3.0 | 0.6 |
| CB | M2(P) | 0.0 | 0.7 | 0.6 |
| AT | S1-ul/ll | 0.0 | 1.8 | 0.6 |
| NT | S1-nose | 0.0 | 3.0 | 0.6 |
| YT | M2(P)/RSP | −1.2 | 0.7 | 0.6 |
| RT | S1-ll | −1.2 | 1.8 | 0.6 |
| CT | S1-bfd | −1.2 | 3.0 | 0.6 |
| NA | RSP | −2.4 | 0.7 | 0.6 |
| YA | V2M/PtA | −2.4 | 1.8 | 0.6 |
| RA | V1(A) | −2.4 | 3.0 | 0.6 |
| CA | AuD/A1 | −2.4 | 4.2 | 0.6 |
| YN | RSP/V2M | −3.6 | 0.7 | 0.6 |
| NR | V1(P)/V2M | −3.6 | 1.8 | 0.6 |
| CN | V1(P) | −3.6 | 3.0 | 0.6 |
| YR | AuD/TeA | −3.6 | 4.2 | 0.6 |
| CY | V1(P) | −4.5 | 1.8 | 0.6 |
| CR | V1(P)/V2L | −4.5 | 3.0 | 0.6 |
| *Cortex-wide injections (Figures 2 and 5)* |  |  |  |  |
| AB | mPFC (PrL) | +2.0 | 0.3 | 1.5 |
| BY | M1(A) | +2.0 | 2.0 | 0.6 |
| CB | M1(P) | +1.0 | 1.5 | 0.6 |
| BN | M2(A) | +2.0 | 0.8 | 0.6 |
| RB | M2(P) | +0.5 | 0.5 | 0.6 |
| NA | S1(FL) | 0.0 | 2.5 | 0.6 |
| YA | S1(BF) | −1.0 | 3.0 | 0.6 |
| YR | S2 | −1.0 | 4.0 | 0.6 |
| RA | RSP | −2.0 | 0.8 | 0.6 |
| NR | V2L | −3.0 | 2.5 | 0.6 |
| CN | V1(P) | −4.0 | 3.0 | 0.6 |
| CA | V2M | −2.5 | 1.5 | 0.6 |
| YN | V1(A) | −2.5 | 3.2 | 0.6 |
| CY | A1 | −2.5 | 4.5 | 0.6 |
| *Motor and somatosensory injections (Figure 3)* |  |  |  |  |
| NT | M2 (face) | +2.1 | 0.8 | 0.6 |
| RT | M2 (body) | +1.2 | 0.5 | 0.6 |
| YR | M1 (face) | +1.5 | 2.2 | 0.6 |
| YT | M1-ul | +0.5 | 1.2 | 0.6 |
| YN | M1-ll | −0.2 | 1.0 | 0.6 |
| CR | S1-mouth | +0.85 | 3.3 | 0.6 |
| CN | S1-nose | +0.4 | 2.8 | 0.6 |
| CT | S1-ul | 0.0 | 2.2 | 0.6 |
| CY | S1-ll | −0.5 | 1.6 | 0.6 |
| NR | S1-bfd | −1.0 | 3.0 | 0.6 |
| *Marmoset V1 injections (Figure 6 and 7)* |  |  |  |  |
| CR | V1 (exposed) | −12.46 | 10.15 | 0.8 |
| YR | V1 (exposed) | −12.44 | 8.75 | 0.8 |
| RA | V1 (exposed) | −12.44 | 6.83 | 0.8 |
| RT | V1 (exposed) | −12.44 | 4.51 | 0.8 |
| CT | V1 (exposed) | −12.44 | 1.96 | 0.8 |
| YT | V1 (exposed) | −14.20 | 2.13 | 0.8 |
| YN | V1 (exposed) | −14.20 | 4.90 | 0.8 |
| CY | V1 (exposed) | −14.20 | 7.11 | 0.8 |
| CN | V1 (exposed) | −14.20 | 8.41 | 0.8 |
| NT | V1 (exposed) | −15.56 | 4.73 | 0.8 |
| NA | V1 (buried) | −7.63 | 3.14 | 3.9 |
| YA | V1 (buried) | −10.41 | 3.14 | 4.0 |
| NR | V1 (buried) | −12.53 | 3.14 | 3.9 |
| AT | V1 (buried) | −12.65 | 5.91 | 6.4 |
| CA | V1 (buried) | −14.22 | 4.04 | 2.9 |

**Table S2.** Allen Mouse Brain Connectivity Atlas experiments corresponding to FluoCode injection sites and correlation of their thalamic projection patterns. For each of the 14 FluoCode injection sites, a corresponding wild-type experiment was manually selected from the Allen Mouse Brain Connectivity Atlas (Cre driver lines excluded), based on the similarity in coordinate information. Spearman’s rank correlation coefficients (*r*) between FluoCode and Allen projection densities were calculated across 34 thalamic regions for each matched pair. *p* values were adjusted for 14 comparisons by the Holm method. (A) and (P) denote anterior and posterior parts. Other abbreviations are as in Table S3.

| **FluoCode** | **Cortical area** | **Allen experiment ID** | **Spearman *r*** | ***p* (Holm)** |
|---|---|---|---|---|
| AB | mPFC | 157711748 | 0.892 | 1.7 × 10<sup>−11</sup> |
| BY | M1(A) | 584903636 | 0.912 | 8.2 × 10<sup>−13</sup> |
| CB | M1(P) | 127084296 | 0.651 | 9.2 × 10<sup>−5</sup> |
| BN | M2(A) | 112952510 | 0.869 | 2.7 × 10<sup>−10</sup> |
| RB | M2(P) | 141603190 | 0.466 | 5.5 × 10<sup>−3</sup> |
| NA | S1(FL) | 180718587 | 0.81 | 5.2 × 10<sup>−8</sup> |
| YA | S1(BF) | 112951804 | 0.732 | 3.5 × 10<sup>−6</sup> |
| YR | S2 | 117298988 | 0.744 | 2.3 × 10<sup>−6</sup> |
| RA | RSP | 100140949 | 0.607 | 2.8 × 10<sup>−4</sup> |
| NR | V1(A) | 277713580 | 0.829 | 1.3 × 10<sup>−8</sup> |
| CN | V1(P) | 309003780 | 0.87 | 2.6 × 10<sup>−10</sup> |
| CA | V2M | 126861679 | 0.759 | 1.2 × 10<sup>−6</sup> |
| YN | V2L | 100141473 | 0.934 | 1.0 × 10<sup>−14</sup> |
| CY | A1 | 116903230 | 0.763 | 1.1 × 10<sup>−6</sup> |

**Table S3.** Abbreviations of cortical injection sites and target brain regions used in Figure 5 and Figure S6. Abbreviations of the cortical injection sites and of the thalamic and non-thalamic subcortical regions used in Figures 5 and S6. The region names and hierarchical groupings adhere to the Allen Mouse Brain Common Coordinate Framework (CCFv3), unless otherwise noted. Thalamic nuclei are grouped into medial, intralaminar, midline, ventral, anterior, geniculate, lateral, and reticular subdivisions, and non-thalamic regions are grouped by major division (CTXsp, cortical subplate; CNU, cerebral nuclei; HY, hypothalamus; MB, midbrain; P, pons). Where an abbreviation used in the figures differs from the CCFv3 acronym, the CCFv3 acronym is also listed. Regions shown as single structures in Figure 5D (SPF, MGN, and LGN) are subdivided in Figure S6 (SPFm/SPFp; MGd/MGv/MGm; LGd/LGv/IGL). For cortical areas with two injection sites, (A) and (P) denote the anterior and posterior sites, respectively.

| Abbreviation (figure) | Full name | Category | Group | Allen CCFv3 acronym |
|---|---|---|---|---|
| mPFC | Medial prefrontal cortex | Cortical (injection site) |  |  |
| M1(A) | Primary motor cortex, anterior part | Cortical (injection site) |  | MOp |
| M1(P) | Primary motor cortex, posterior part | Cortical (injection site) |  | MOp |
| M2(A) | Secondary motor cortex, anterior part | Cortical (injection site) |  | MOs |
| M2(P) | Secondary motor cortex, posterior part | Cortical (injection site) |  | MOs |
| S1(FL) | Primary somatosensory cortex, forelimb region | Cortical (injection site) |  | SSp-ul |
| S1(BF) | Primary somatosensory cortex, barrel field | Cortical (injection site) |  | SSp-bfd |
| S2 | Secondary somatosensory cortex | Cortical (injection site) |  | SSs |
| RSP | Retrosplenial agranular cortex | Cortical (injection site) |  | RSPagl |
| V1(A) | Primary visual cortex, anterior part | Cortical (injection site) |  | VISp |
| V1(P) | Primary visual cortex, posterior part | Cortical (injection site) |  | VISp |
| V2M | Secondary visual cortex, medial part | Cortical (injection site) |  |  |
| V2L | Secondary visual cortex, lateral part | Cortical (injection site) |  |  |
| A1 | Primary auditory cortex | Cortical (injection site) |  | AUDp |
| MD | Mediodorsal nucleus of the thalamus | Thalamic | Medial | MD |
| IMD | Intermediodorsal nucleus of the thalamus | Thalamic | Medial | IMD |
| Sm | Submedial nucleus of the thalamus | Thalamic | Medial | SMT |
| CM | Central medial nucleus of the thalamus | Thalamic | Intralaminar | CM |
| CL | Central lateral nucleus of the thalamus | Thalamic | Intralaminar | CL |
| PCN | Paracentral nucleus | Thalamic | Intralaminar | PCN |
| Pf | Parafascicular nucleus | Thalamic | Intralaminar | PF |
| Re | Nucleus of reuniens | Thalamic | Midline | RE |
| Rh | Rhomboid nucleus | Thalamic | Midline | RH |
| PVT | Paraventricular nucleus of the thalamus | Thalamic | Midline | PVT |
| VPM | Ventral posteromedial nucleus of the thalamus | Thalamic | Ventral | VPM |
| VPL | Ventral posterolateral nucleus of the thalamus | Thalamic | Ventral | VPL |
| VM | Ventral medial nucleus of the thalamus | Thalamic | Ventral | VM |
| VAL | Ventral anterior-lateral complex of the thalamus | Thalamic | Ventral | VAL |
| SPF | Subparafascicular nucleus | Thalamic | Ventral | SPF |
| SPFm | Subparafascicular nucleus, magnocellular part | Thalamic | Ventral | SPFm |
| SPFp | Subparafascicular nucleus, parvicellular part | Thalamic | Ventral | SPFp |
| LD | Lateral dorsal nucleus of the thalamus | Thalamic | Anterior | LD |
| AM | Anteromedial nucleus of the thalamus | Thalamic | Anterior | AM |
| AV | Anteroventral nucleus of the thalamus | Thalamic | Anterior | AV |
| AD | Anterodorsal nucleus of the thalamus | Thalamic | Anterior | AD |
| IAD | Interanterodorsal nucleus of the thalamus | Thalamic | Anterior | IAD |
| MGN | Medial geniculate nucleus | Thalamic | Geniculate | MG |
| MGd | Medial geniculate complex, dorsal part | Thalamic | Geniculate | MGd |
| MGv | Medial geniculate complex, ventral part | Thalamic | Geniculate | MGv |
| MGm | Medial geniculate complex, medial part | Thalamic | Geniculate | MGm |
| LGN | Lateral geniculate nucleus | Thalamic | Geniculate | LGd + LGv + IGL |
| LGd | Lateral geniculate nucleus, dorsal part | Thalamic | Geniculate | LGd |
| LGv | Lateral geniculate nucleus, ventral part | Thalamic | Geniculate | LGv |
| IGL | Intergeniculate leaflet of the lateral geniculate complex | Thalamic | Geniculate | IGL |
| LP | Lateral posterior nucleus of the thalamus | Thalamic | Lateral | LP |
| POm | Posterior medial nucleus of the thalamus | Thalamic | Lateral | PO |
| POL | Posterior limiting nucleus of the thalamus | Thalamic | Lateral | POL |
| Eth | Ethmoid nucleus of the thalamus | Thalamic | Lateral | Eth |
| PoT | Posterior triangular thalamic nucleus | Thalamic | Lateral | PoT |
| SGN | Suprageniculate nucleus | Thalamic | Lateral | SGN |
| TRN | Thalamic reticular nucleus | Thalamic | Reticular | RT |
| CLA | Claustrum | Non-thalamic | CTXsp | CLA |
| EP | Endopiriform nucleus | Non-thalamic | CTXsp | EP |
| CP | Caudoputamen | Non-thalamic | CNU | CP |
| ACB | Nucleus accumbens | Non-thalamic | CNU | ACB |
| FS | Fundus of striatum | Non-thalamic | CNU | FS |
| GPe | Globus pallidus, external segment | Non-thalamic | CNU | GPe |
| GPi | Globus pallidus, internal segment | Non-thalamic | CNU | GPi |
| SI | Substantia innominata | Non-thalamic | CNU | SI |
| LA | Lateral amygdalar nucleus | Non-thalamic | CNU | LA |
| BLAa | Basolateral amygdalar nucleus, anterior part | Non-thalamic | CNU | BLAa |
| MEA | Medial amygdalar nucleus | Non-thalamic | CNU | MEA |
| ZI | Zona incerta | Non-thalamic | HY | ZI |
| LHA | Lateral hypothalamic area | Non-thalamic | HY | LHA |
| SCs | Superior colliculus, sensory related | Non-thalamic | MB | SCs |
| SCm | Superior colliculus, motor related | Non-thalamic | MB | SCm |
| IC | Inferior colliculus | Non-thalamic | MB | IC |
| APN | Anterior pretectal nucleus | Non-thalamic | MB | APN |
| NOT | Nucleus of the optic tract | Non-thalamic | MB | NOT |
| OP | Olivary pretectal nucleus | Non-thalamic | MB | OP |
| PPT | Posterior pretectal nucleus | Non-thalamic | MB | PPT |
| SNr | Substantia nigra, reticular part | Non-thalamic | MB | SNr |
| SNc | Substantia nigra, compact part | Non-thalamic | MB | SNc |
| RN | Red nucleus | Non-thalamic | MB | RN |
| PAG | Periaqueductal gray | Non-thalamic | MB | PAG |
| PG | Pontine gray | Non-thalamic | P | PG |
| CTXsp | Cortical subplate | Major division |  | CTXsp |
| CNU | Cerebral nuclei | Major division |  | CNU |
| HY | Hypothalamus | Major division |  | HY |
| MB | Midbrain | Major division |  | MB |
| P | Pons | Major division |  | P |

**Table S4.**
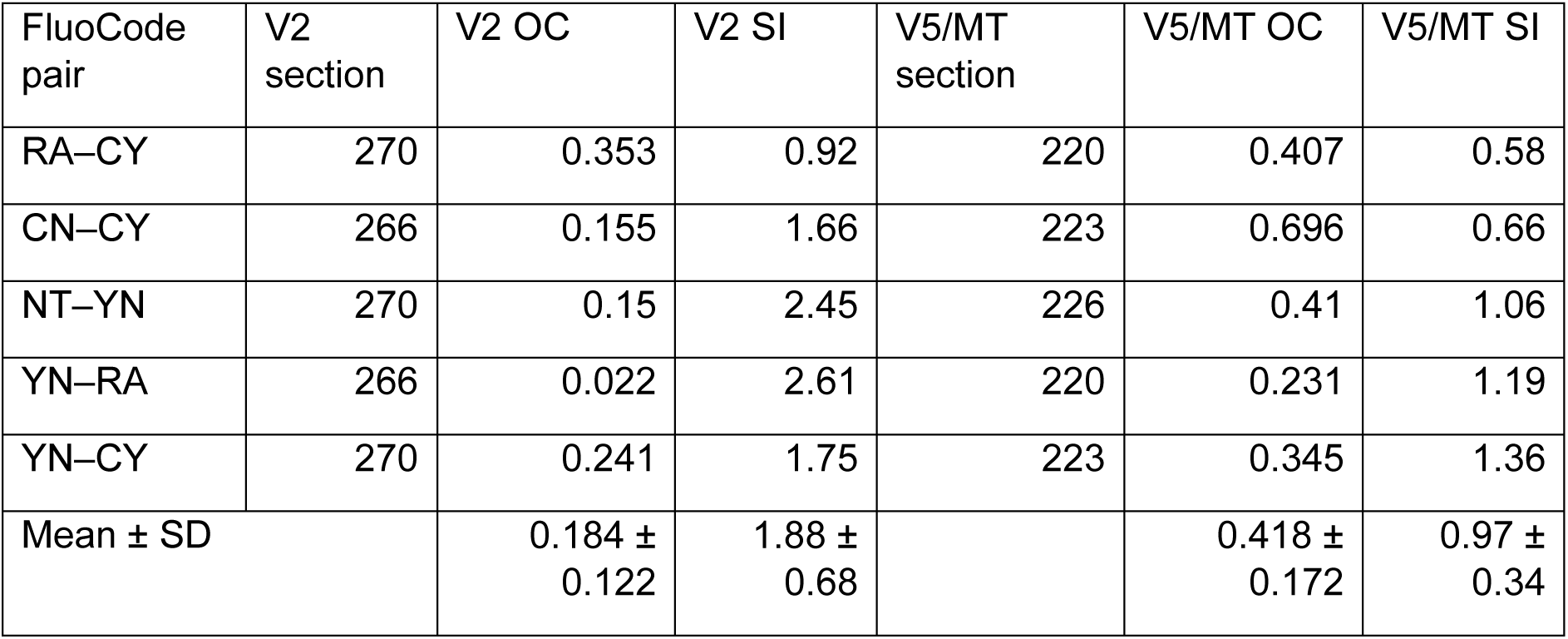
Segregation of adjacent FluoCode terminal fields in V2 and V5/MT. Overlap coefficient (OC) and separation index (SI) were calculated from the normalized intensity profiles of each FluoCode pair, measured along the cortical surface with a 300-µm-wide ROI (see Methods). Lower OC and higher SI indicate greater segregation. Section, the section in which each pair was analyzed. The bottom row shows mean ± SD across the five pairs. V2 vs V5/MT: OC, p = 0.052; SI, p = 0.018 (two-tailed paired t-test, n = 5 FluoCode pairs).

| FluoCode pair | V2 section | V2 OC | V2 SI | V5/MT section | V5/MT OC | V5/MT SI |
|---|---|---|---|---|---|---|
| RA–CY | 270 | 0.353 | 0.92 | 220 | 0.407 | 0.58 |
| CN–CY | 266 | 0.155 | 1.66 | 223 | 0.696 | 0.66 |
| NT–YN | 270 | 0.15 | 2.45 | 226 | 0.41 | 1.06 |
| YN–RA | 266 | 0.022 | 2.61 | 220 | 0.231 | 1.19 |
| YN–CY | 270 | 0.241 | 1.75 | 223 | 0.345 | 1.36 |
| Mean ± SD |  | 0.184 ± 0.122 | 1.88 ± 0.68 |  | 0.418 ± 0.172 | 0.97 ± 0.34 |

### Supplementary Videos

**Supplementary Video 1. FASTd serial sections across the mouse brain.**

**Supplementary Video 2. 3D rendering of a whole mouse brain image obtained with FASTd.**

**Supplementary Video 3. 3D rendering of the FluoCode signals within the individual thalamic nuclei the mouse.**

**Supplementary Video 4. Descending projections from cortical areas in mice visualized with light-sheet microscopy.**

**Supplementary Video 5. Serial mouse brain slices analyzed with the FluoCode Reader.**

**Supplementary Video 6. Serial sections of a marmoset obtained with STPT.**

**Supplementary Video 7. 3D rendering of a whole marmoset brain obtained with STPT.**

